# SpaCeNet: Spatial Cellular Networks from omics data

**DOI:** 10.1101/2022.09.01.506219

**Authors:** Stefan Schrod, Niklas Lück, Robert Lohmayer, Stefan Solbrig, Dennis Völkl, Tina Wipfler, Katherine H. Shutta, Marouen Ben Guebila, Andreas Schäfer, Tim Beißbarth, Helena U. Zacharias, Peter J. Oefner, John Quackenbush, Michael Altenbuchinger

## Abstract

Advances in omics technologies have allowed spatially resolved molecular profiling of single cells, providing a window not only into the diversity and distribution of cell types within a tissue, but also into the effects of interactions between cells in shaping the transcriptional landscape. Cells send chemical and mechanical signals which are received by other cells, where they can subsequently initiate context-specific gene regulatory responses. These interactions and their responses shape the individual molecular phenotype of a cell in a given microenvironment. RNAs or proteins measured in individual cells together with the cells’ spatial distribution provide invaluable information about these mechanisms and the regulation of genes beyond processes occurring independently in each individual cell. “SpaCeNet” is a method designed to elucidate both the intracellular molecular networks (how molecular variables affect each other within the cell) and the intercellular molecular networks (how cells affect molecular variables in their neighbors). This is achieved by estimating conditional independence relations between captured variables within individual cells and by disentangling these from conditional independence relations between variables of different cells. A python implementation of SpaCeNet is publicly available at https://github.com/sschrod/SpaCeNet.

## 1 Introduction

Measurements of spatially resolved RNA or protein expression patterns open unprecedented opportunities to study questions in areas such as developmental biology and pathophysiology, where interactions between cells are known to influence a wide range of processes. Methods such as “spatial transcriptomics” [1] and Slide-Seq [2] used molecular barcoding to count mRNAs aggregated in small regions and integrated those with images to produce a transcript map. A drawback of these early methods was that mRNAs from multiple cells in a small region could contribute to the observed signal, masking differences between cells and cell-cell interactions. Subsequent barcoding techniques substantially improved resolution, such as Slide-Seq [3] and Slide-SeqV2 [4] with a resolution of *∼* 10μm, as well as, most recently, Seq-Scope [5] and Stereo-Seq [6] with a resolution of *∼* 0.5μm. *In situ* hybridization or sequencing methods can measure the expression of many genes with single-cell resolution (and even subcellular resolution) [7, 8, 9, 10], but these methods require complex instrumentation and long imaging times. Three-dimensional intact tissue sequencing in single cells has been achieved by STARmap (spatially-resolved transcript amplicon readout mapping) [11], which is capable of measuring simultaneously the expression of about 1000 genes at single-cell resolution in a three-dimensional thick tissue section. Although spatial omics technologies are in their infancy, the launch of commercial products such as the 10X Genomics Visium platform has led to increased interest in methods that will allow the analysis, interpretation and exploration of data generated [12, 13, 14, 15, 16, 17].

Cells organize themselves spatially within tissues and organisms in order to carry out specific functions. This organization is orchestrated via signals that include physical interactions via cell-cell contact, chemical signals, and exosome-mediated transfer of RNAs between cells. Each cell’s individual phenotype together with its location in space relative to other cells captures information about this process. For instance, genes encoding chemokines (chemoattractant cytokines that facilitate intercellular communication) are first transcribed and then translated before the respective proteins pass through the cell membrane into the extracellular domain where they recruit leukocytes. When these signals are received by the leukocytes, they initiate signaling cascades that finally induce a molecular response leading the cells to adapt their individual molecular phenotype and alter their behavior. While such interactions are known to be essential for many biological processes [18], there are no well-established statistical methods to investigate the relationships between spatial organization, gene or protein expression, and cellular phenotype. There are first approaches to infer cellular interactions from spatial omics, although they differ substantially with respect to semantics, modelling approaches, and overall objectives. In seminal work [13], Arnol et al. proposed Spatial Variance Component Analysis (SVCA) to decompose the expression of individual genes into spatial and non-spatial contributions, namely into cell intrinsic effects, general environmental effects, and cell-cell interaction effects. SVCA is motivated by Gaussian processes and models gene/protein expression independently of each other. Thus, it does not capture complex multivariate relationships between genes [13], but detects individual genes which are involved in cell-cell interactions. Another approach is Node-Centric Expression Modeling (NECM) [19]. NECM uses a graph neural network to predict cells’ observed gene expression vector from respective cell type label and niche, with the latter resembling cell-cell communication in terms of statistical dependencies between cells. As such, NECM identifies cell-type couplings. However, it does not identify gene pairs involved in cellular interactions. This was addressed by COMMunication analysis by Optimal Transport (COMMOT) using the framework of optimal transport to account for the competition among ligand and receptor species, while taking into account the spatial distances between cells [20], by Mixture of Experts for Spatial Signaling genes Identification (MESSI) considering an additional cell neighbourhood to infer intra-cellular responses [21], and by Multiview Intercellular SpaTial modeling framework (MISTY) using an aggregated direct neighbourhood to resolve spatial effects [22]. However, rigorous statistical models to infer complex cellular interdependencies from spatially distributed molecular data are still missing.

The statistical inference of correlation-based molecular networks from high-dimensional omics data is based on the assumption that coordinated expression in a cell might provide insight into processes that are activated or inactivated in different phenotypes. Early attempts used pairwise measures of gene-gene coexpression such as mutual exclusivity, Pearson’s and Spearman’s correlations and identified network edges between genes based on a correlation threshold [23, 24]. Such measures of coexpression can provide insight into active biological mechanisms, but they are vulnerable to identifying spurious associations [25]. These associations can be the consequence of indirect dependencies that cannot be resolved if pairwise relationships between molecular variables are considered only in isolation from all other molecular variables. This stimulated research in high-dimensional statistics, specifically in Probabilistic Graphical Models (PGMs). PGMs resolve the dependency structure of molecular variables and, thus, disentangle direct from indirect associations. This is even possible in high-dimensional settings, where the number of variables is larger than the number of observations. The main concept underlying PGMs is conditional independence. Two variables *X* and *Y* are considered as conditionally independent (given all remaining variables), if *X* does not provide any additional information about *Y* that is not already covered by the remaining variables. Thus, although *X* and *Y* might be pairwise correlated, they can still be conditionally independent; the pairwise correlation could be just the consequence of *X*’s and *Y* ‘s indirect relationship mediated via other variables and not due to a direct relationship [26]. This powerful concept made PGMs one of the favored approaches to resolve molecular networks from molecular data [24] and, importantly, they can be straightforwardly extended to account for complex single-cell and multi-omics data [27, 28]. It is nevertheless noteworthy that PGMs are undirected graphical models and do not resolve causal relationships, although, intriguingly, lower bounds on causal effects can be provided [29].

Spatial Cellular Networks from omics data (SpaCeNet) is a method for analyzing patterns of correlation in spatial transcriptomics data by extending the concept of conditional independence to spatially distributed information, facilitating reconstruction of both the intracellular and the intercellular interaction networks with single-cell spatial resolution. SpaCeNet was developed to address the diversity of cellular interactions and the various length scales over which they occur. SpaCeNet introduces flexible interaction potentials in combination with appropriate regularization strategies to allow this diversity in cellular state, tissue organisation and spatial communication to be handled effectively. We validate SpaCeNet in extensive simulation studies and illustrate its capacity to augment exploratory data analysis of spatial transcriptomics data from the mouse visual cortex and the *Drosophila* blastoderm.

## 2 Results

### 2.1 SpaCeNet infers cell-cell interactions from spatial omics data

#### SpaCeNet concept

The individual molecular phenotype of a cell is shaped by its microenvironment and the underlying processes are diverse, comprising the exchange of signals via direct physical contact or via signalling molecules. Throughout the article, we will refer to any process with which cells affect each others molecular appearance simply as cell-cell or cellular “interactions”. Spatial Transcriptomics (ST) involves measuring cellular phenotypes along with cells’ location in space, providing information about such processes and serving as a valuable resource to study cell-cell interactions. To develop SpaCeNet, we require single-cell resolved ST data (single-cell profiles + cellular positions) such that cells can be described as interacting units. In recent years, such data have become increasingly available via both multiplexing and barcoding techniques [11, 5, 6].

Key to SpaCeNet is a statistically sound approach that uses probabilistic graphical models (PGMs) to decompose the observed cellular profiles into contributions arising from ordinary cellular variability and contributions from cellular interactions. Formally, SpaCeNet decomposes each cell’s observed molecular profile into (i) a baseline contribution corresponding to the cell’s molecular profile in isolation (Figure 1A) and (ii) a residual contribution attributed to its interaction with other cells (Figure 1B and C). Importantly, this decomposition uses a model parametrization such that vanishing parameters encode conditional independence statements, namely intracellular and intercellular spatial conditional independence (SCI) relationships as introduced in the following and as summarized in Suppl. Figure S1.

**Figure 1:**
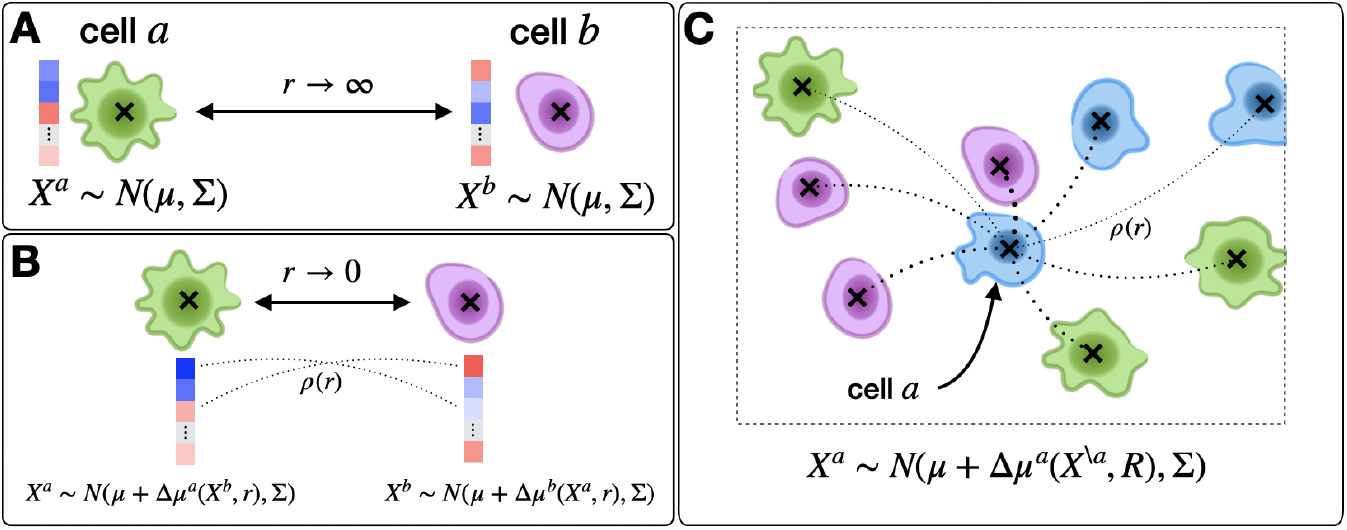
SpaCeNet concept and how it applies to different cellular contexts. Figure **A** shows a schematic picture of two isolated cells *a* and *b* which are infinitely separated (distance *r → ∞*). In this case, respective cellular profiles with expression levels illustrated in blue and red are modeled by a multivariate normal distribution with mean expression *μ* and covariance Σ. The latter encodes the complex co-expression pattern of different RNAs/proteins and facilitates a description of molecularly diverse cells. Figure **B** shows for comparison a scenario of two neighboring cells, where the cells affect each other; the molecular phenotype changes compared to Figure **A** as a consequence of the different cellular context. This molecular adaptation of cell *a* (analogously for cell *b*) is modeled by a shifted mean expression vector *μ*+Δ*μ*^*a*^(*X*^*b*^, *r*), where the molecular adaptation depends on the molecular phenotype of cell *b* and the cell-cell distance *r*. The molecular adaptation is parametrized by interaction potentials which directly provide estimates of spatial gene-gene dependencies between gene *i* and *j*. Figure **C** sketches the more complex scenario of a set of interacting cells. As such, the expression of cell *a* is affected by all surrounding cells in a distance-dependent way, as illustrated by thin and bold dotted lines for long- and short-range interactions, respectively. One should note that the mean expression of cell *a* now depends on the molecular phenotype of all other cells *X*^*\a*^ as well as all respective cell-cell distances summarized in *R*.

PGMs are graphical models in which nodes represent variables assumed to be distributed according to a probabilistic model and edges between nodes represent conditional dependence between them, where the conditioning is on the remaining nodes in the network. The absence of edges in a PGM defines a set of Conditional Independence (CI) relations corresponding to direct pairwise independence between variables: for two variables *X* and *Y* that are CI, any association between *X* and *Y* observed in the data can be attributed to indirect effects of other variables in the system. Conversely, if *X* and *Y* are not CI, i.e., there is an edge between the two, this dependence is not mediated by other variables in the system. As such, PGMs identify and remove potential erroneous (false-positive; indirect) associations. This makes PGMs versatile tools to infer gene-association networks, as repeatedly shown in the literature [24, 25].

Here, we develop a similar framework to model intracellular and intercellular CI relations in ST data. We estimate SCI between variable *X*^*a*^ (gene *X* in cell *a*) and variable *Y* ^*b*^ (gene *Y* in a neighboring cell *b*) keeping all other variables in the data fixed; similar to CI, by conditioning on all variables except for *X*^*a*^ and *Y* ^*b*^, we can study the direct dependence of *X*^*a*^ on *Y* ^*b*^ and vice versa. As such, SpaCeNet disentangles (1) direct from indirect relationships of variables in individual cells, and (2) direct from indirect relationships between molecular variables in different cells. A direct consequence of both is that each cell’s environmental adaptation in its spatial context can be estimated distinctly from its molecular phenotype in isolation. Since we are interested in sample estimates of cellular interaction patterns, SpaCeNet infers SCI simultaneously across all pairs of cells in a ST dataset.

#### SpaCeNet infers both short and long-range cell-cell interactions while taking into account the molecular diversity of cells

SpaCeNet encodes intercellular SCI relationships via interaction potentials mediating the association strength between molecular variables across cells in a distance dependent manner. We denote the interaction potential between *X*^*a*^ (gene *X* in cell *a*) and *Y* ^*b*^ (gene *Y* in cell *b*) at distance *r*_*ab*_ as *ρ*_*XY*_ (*r*_*ab*_). Since *ρ*_*XY*_ (*r*_*ab*_) depends only on the distance *r*_*ab*_ between two cells, the intercellular potential between genes *X* and *Y* takes the same functional form across all modeled cells and, moreover, vanishing potentials *ρ*_*XY*_ (*r*_*ab*_) imply that *X*^*a*^ and *Y* ^*b*^ are SCI across all cells *a* and *b*. One might ask whether such a one-function-fits-all approach is appropriate to model complex cellular communication between molecularly diverse cells, considering that even related cell types, such as T-cells and B-cells, fulfill very different tasks and are expected to send and receive very different signals. As briefly illustrated, SpaCeNet can encode such diverse cell-cell communication patterns taking into account the individual cell’s molecular phenotype without requiring *a priori* specification of involved cell types, variables, and interaction mechanisms. Assuming all variables except *X*^*a*^ and *Y* ^*b*^ to be fixed, *X*^*a*^ depends on *Y* ^*b*^ via the regression formula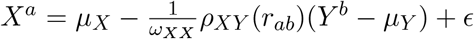 (this can be also seen by marginalizing the full density Eq. (1) over all variables except *X*^*a*^ and *Y* ^*b*^). Thus, the effect that molecular variable *Y* ^*b*^ in cell *b* exerts on *X*^*a*^ in cell *a* via cell-cell interaction is directly related to the individual molecular profile of cell *b*, which means that the molecular phenotype of a cell determines how it communicates with other cells. One should also note that SpaCeNet infers the *ρ*_*XY*_ (*r*) without assuming a particular functional dependence on *r*_*ab*_ via series expansion in powers of 1*/r*_*ab*_. This series expansion is motivated by the fact that infinitely separated cells cannot communicate through the release or absorbance of particular signaling molecules and the drop in concentration of these molecules with distance suggests that *ρ*_*XY*_ (*r*_*ab*_) should vanish for *r*_*ab*_ *→ ∞*.

To show that SpaCeNet can reconstruct diverse cell-cell interactions, we performed four illustrative simulations shown in Figure 2a to d, where we simulated radial dependencies corresponding to long-distance interactions (such as via paracrine signaling, Figure 2a and b), short-distance interactions (such as via cell-cell contact) using an exponentially decreasing potential with short range (Figure 2c), and an interaction where the potential first grows with *r*_*ab*_, peaks at average distances, and then goes to zero for large *r*_*ab*_ (Figure 2d). The precise radial dependencies are provided in the caption of Figure 2. In principle, SpaCeNet can model interaction potentials to arbitrary orders in 1*/r*_*ab*_. We present the corresponding radial dependencies estimated via SpaCeNet for expansions up to order (1*/r*_*ab*_)^*L*^ for *L* = 1 (solid lines), *L* = 3 (dashed lines), and *L* = 10 (dotted lines). As expected, the approximation of the true underlying interaction improves with increasing *L*. However, increasing the order in 1*/r*_*ab*_ results in higher model complexity, which in turn comes at the cost of additional parameters, increasing the risk of overfitting and computational burden.

**Figure 2:**
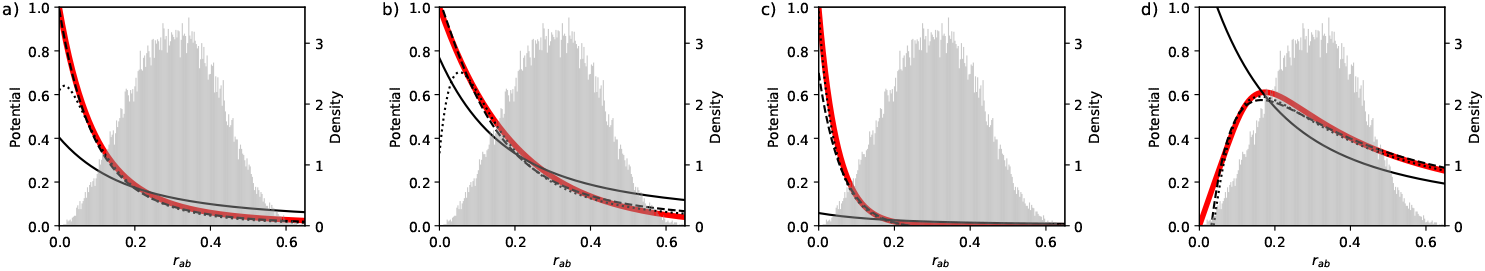
SpaCeNet can reconstruct diverse cellular interaction patterns. Figures a to d show different potentials used to simulate data in red in dependence of the distance between cells *r*_*ab*_. Potentials inferred by SpaCeNet are shown for three different expansion orders: *L* = 1 (black solid lines), *L* = 3 (black dashed lines), and *L* = 10 (black dotted lines), illustrating that with higher order, increasingly complex cellular interactions can be reconstructed. The corresponding ground-truth potentials (red lines) are (a) a standard decreasing potential corresponding to the second-order term in the series expansion, *ρ*(*r*_*ab*_) = (1 *−* exp(*−r*_*ab*_*/r*_0_))^2^(*r*_0_*/r*_*ab*_)^2^ with *r*_0_ = 1*/*10, (b) a long-range exponential potential *ρ*(*r*_*ab*_) = exp(*−*5 *r*_*ab*_), (c) a short-range exponential potential *ρ*(*r*_*ab*_) = exp(*−*20 *r*_*ab*_), and (d) a potential *ρ*(*r*_*ab*_) = 10 (1 *−* exp(*−r*_*ab*_*/*2)) (1 *−* exp(*−*1*/*(5*r*_*ab*_)^2^)) which increases for small *r*_*ab*_ and then decreases again. The corresponding densities of the pairwise distances *r*_*ab*_ are shown as histograms in the background of the figures with the respective *y*-axis depicted at the figures’ right axis. For more details about data simulation see Suppl. Sect. 1.

#### SpaCeNet recovers inter-and intracellular interactions from ***in silico*** generated tissues

We systematically evaluated SpaCeNet’s capacity to reconstruct intra- and intercellular networks defined in terms of SCI relationships. To this end, we generated *in silico* tissues from the full probability density Eq. (1) and tested if the ground truth intra- and intercellular network edges are correctly recovered (see also Suppl. Section 2). Since cellular interactions occur at various length scales, we performed three different studies involving (1) long-range interactions, (2) medium-range interactions, (3) short-range interactions, and (4) a mixture of long-, medium- and short-range interactions. For each study, we simulated a total of *n · S ∈ {*10^3^, 10^4^, 10^5^*}* cells, where *n* is the number of cells per ST slide and *S ∈ {*1, 10, 100*}* the number of ST slides. Performance was assessed using the area under the precision recall curve (AUPRC) and the area under the receiver operating characteristic curve (AUROC) (see Suppl. Table S1).

In our first simulation study we used an exponentially decreasing interaction potential *ρ*_*ij*_(*r*_*ab*_) = Δ*ρ*_*ij*_ *·* exp(*−ϕ*_*ij*_*r*_*ab*_) with a comparatively large range of 1*/ϕ*_*ij*_ = 1*/*5. Suppl. Figure S4A gives AUPRC versus *n · S* to recover the correct intracellular networks (the intracellular precision matrix **Ω**, left figure) and the intercellular interaction networks (the cell-cell interaction parameters **Δ*ρ***, right figure). We find that SpaCeNet is capable of reconstructing intracellular networks across the full range of simulated cells, *n · S*, with a median AUROC larger than *∼* 0.85 throughout all settings. Edge recovery of cell-cell interactions was achieved with a median AUPRC ranging from 0.28 for *n* = 10^3^ and *S* = 1 (blue) to 0.92 for *n* = 10^3^ and *S* = 100. It is worth noting that edge recovery for both the intra- and intercellular networks was slightly better if cells are distributed across measurements, as seen by comparison of the AUPRC for *n* = 10^3^ and *S* = 1 (blue), for *n* = 100 and *S* = 10 (orange), and for *n* = 10 and *S* = 100 (green), that all share the same total number of cells *n · S* = 10^3^. This trend also persisted for *n · S* = 10^4^. We also tested the same class of potentials with medium range 1*/ϕ*_*ij*_ = 1*/*10 (Suppl. Figure S4B) and short range 1*/ϕ*_*ij*_ = 1*/*20 (Suppl. Figure S4C). The results were similar to the first simulation study. The edge recovery in terms of AUPRC improved with an increasing number of cells *n · S*, the more so when cells were distributed across multiple samples *S*. Finally, we studied how diverse cell-cell interactions impair model inference (Suppl. Figure S4D). For this, we repeated the previous simulations, but created flexible potentials *ρ*_*ij*_(*r*_*ab*_) = Δ*ρ*_*ij*_ exp(*−ϕ*_*ij*_*r*_*ab*_) with *ϕ*_*ij*_ *∼* Unif(5, 20) and Δ*ρ*_*ij*_ *∼* Unif(*−*1, 1) between molecular variables *i* and *j* that differ with respect to interaction range and strength. For illustration purposes, these different potential ranges are shown in Suppl. Figure S2 for *ϕ*_*ij*_ = 5 (solid line), *ϕ*_*ij*_ = 10 (dashed line), and *ϕ*_*ij*_ = 20 (dotted line). In summary, we found that SpaCeNet achieves an edge-recovery performance that is almost as good as in the more controlled setting of the previous simulation studies, in particular when the total number of cells *n· S* is increased. An exemplary SpaCeNet model obtained in this study together with its ground truth is shown in Suppl. Figure S3 for two different model regularization parameters *β*. Results considering AUROC for all four simulation studies are shown in Suppl. Figure S4, supporting our previous findings.

To further illustrate the relevance of distance dependent interactions, we compared SpaCeNet to two alternative approaches, namely an ordinary spatial correlation and its naive partial extension implemented as follows. For each individual cell *i* and gene *j*, we summed up the expression of the top *K* nearest neighbors, yielding an additional set of “regional expression variables”. Then, we correlated the cell’s expression of gene *j* with the regional expression of gene *k* for all *j* and *k*, yielding a matrix of correlation coefficients. The performance evaluation shown in Suppl. Figure S5 and Suppl. Table S2 contrasts the performance of this naive correlation based approach for *K* = 1, 5, 10, 20, 50 with SpaCeNet. We observe that, first, overall this approach cannot compete with SpaCeNet and, second, that its performance depends on *K*, with long-range interactions typically better captured by large *K* (simulation A) and short range interaction better captured for small *K*. In contrast, SpaCeNet dynamically adjusts for the interaction range and does not require any *a priori* specification of interacting neighbors.

As a further benchmark, we included an extension of the naive spatial correlation measure to a partial correlation measure. Let *j*^*′*^ denote the regional expression variable which represents the expression of gene *j* of the top *K* neighbors. A remedy to estimate spatial associations between genes *j* and *k* has to take into account that both might also correlate to each other within individual cells. Therefore, we decided to calculate partial correlations between variables *j* and *k*^*′*^ conditioning on *j*^*′*^ and *k*. Performance analysis of this additional baseline strongly supports our finding that SpaCeNet, as a dynamic approach that intrinsically infers complex distance dependencies, is much more suited to infer cell-cell interactions (Suppl. Figure S6 and Suppl. Table S3).

#### SpaCeNet recovers intra- and intercellular interactions from ***in silico*** tissues generated via mechanistic modeling

Former simulations illustrate that SpaCeNet can reliably recover SCI relationships. Next, we aimed to verify SpaCeNet in more realistic settings. Tanevski et al. [22] used a comprehensive *in silico* tissue model to mimic the interactions of different cell types through ligand binding and subsequent signaling events. They generated two *in silico* tissues with pre-specified cell-cell interactions between four cell types, capturing interactions of 29 molecular species including 5 ligands, 5 receptors, and 19 intracellular signaling proteins. The authors suggested Multiview Intercellular SpaTial modeling framework (MISTy) to derive importance scores for each pair of markers to infer intracellular and intercellular molecular interactions. We followed their work and considered an intracellular interaction as correct if there is a direct interaction between the markers in the model’s networks and an intercellular interaction as correct if one of the markers is responsible for a ligand production and the other one is activated by it [22]. Respective results in terms of AUPRC are summarized in Table 1 for MISTy and SpaCeNet. To evaluate SpaCeNet’s performance, we restricted ourselves to leading order potentials to ensure comparability. We observe that SpaCeNet consistently outperformed MISTy with respect to both the recovery of intra- and intercellular interactions. This is also supported by a respective analysis considering AUROC (Suppl. Table S4). One should note that MISTy does not disentangle direct from indirect relationships. As a consequence, most false positive interactions derived by MISTy might be the result of indirect or higher-order interactions [22].

**Table 1:**
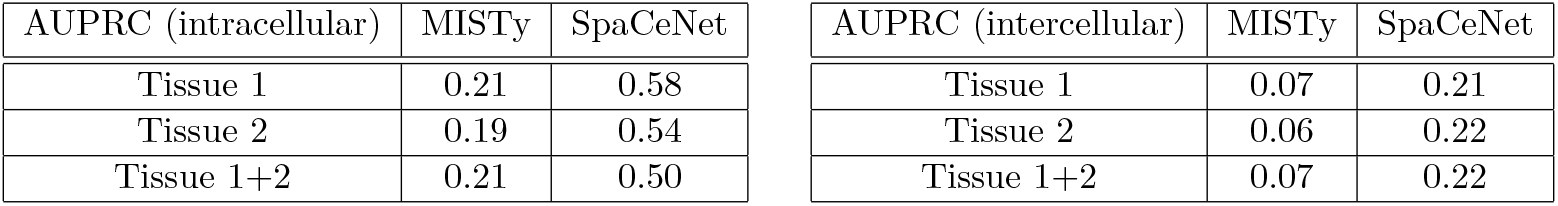
Performance in terms of AUPRC to recover intra-(left) and intercellular (right) interactions from *in silico* tissues generated via mechanistic modeling for MISTy and SpaCeNet.

### 2.2 SpaCeNet resolves molecular mechanisms involved in cellular interactions in the mouse visual cortex

In the following, we present an example analysis using SpaCeNet for spatial transcriptomics data from the mouse visual cortex generated by STARmap [11]. STARmap labels and amplifies cellular RNAs. These amplicons are then transferred to a hydrogel while lipids and proteins are removed. The hydrogel is optically transparent and can be sequentially imaged through multiple cycles with a low probability of errors and miscodings. STARmap measurements of the mouse visual cortex were downloaded from https://www.starmapresources.org/data. The data consist of measurements for 28 RNAs in *∼*30,000 cells together with their respective locations in a 0.1 *×* 1.4 *×* 1.7 mm^3^ tissue section.

We used SpaCeNet to estimate a global, intracellular network (the precision matrix **Ω**) and a network of spatial molecular interactions (the spatial interaction parameters **Δ*ρ***^*(·)*^), where we set *L* = 3 and split the data into four equally sized batches of which three served for model building and one for model validation and hyper-parameter calibration (Suppl. Figure S7). We tuned *α, β ∈* [10^*−*5^, 10] on a 4 *×* 4 grid that was refined 6 times (Suppl. Figure S8) and selected the best set of hyper-parameters based on the highest validation pseudo-log-likelihood. A final model was estimated using the full data set.

For the intracellular precision matrix **Ω** we obtained a complete matrix with weights summarized in Suppl. File 1. For the spatial interactions **Δ*ρ***^*(·)*^, SpaCeNet selected a set of 134 out of 406 possible spatial associations (Suppl. Figure S9, Suppl. File 1). We ranked edges according to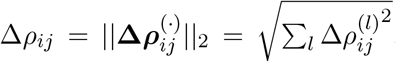, for which the greatest edge weight was between myelin basic protein gene (*Mbp*) and FMS-related receptor tyrosine kinase 1 (*Flt1*) with Δ*ρ*_*Mbp, Flt1*_ = 0.328 and the leading-order contribution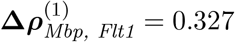. Negative/positive values of 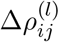 correspond to positive/negative spatial associations in analogy to the definition of the precision matrix **Ω**.

First, we verified the absolute residuals between the observed data matrix **X** and its SpaCeNet reconstruction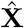 using either (a) the intracellular edges only (**Δ*ρ***^*(·)*^ = **0**), or (b) the full model (see also Suppl. Sect. 2). The results corresponding to this analysis are shown in Figure 3a and b, respectively. We observed that the spatial associations improve model building substantially, as illustrated by the band of large residuals in the upper left quadrant for both *Flt1* and *Mbp* (Figure 3a and d), that were reduced substantially via the spatial interactions (Figure 3b and e). Figure 3c and f show the corresponding contributions from the spatial interactions **Δ*ρ***^*(·)*^.

**Figure 3:**
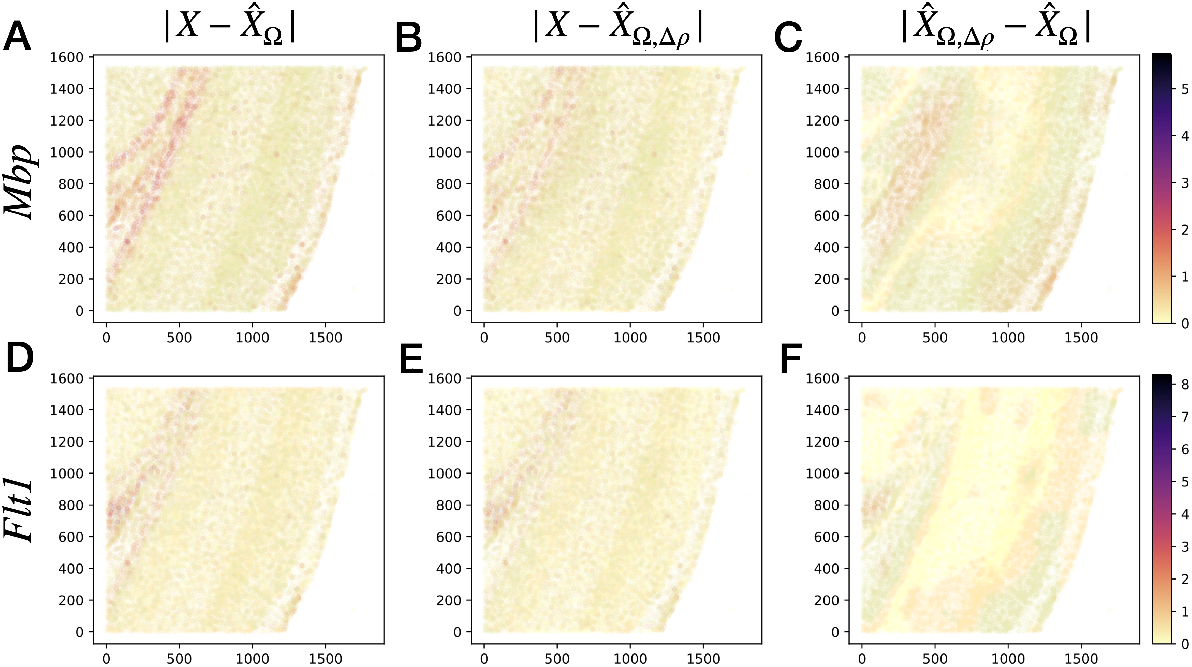
Spatial interactions improve on goodness of fit. The figures show absolute prediction residuals for spatially associated genes *Mbp* (top row) and *Flt1* (bottom row) in the mouse visual cortex data from [11] with respect to their position in space. The left column (Figures A and C) shows the absolute residuals between the ground truth data and the predictions based on the intracellular model parameters only, 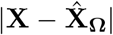The middle column Figures B and E display the residuals if both intra- and intercellular interactions are considered,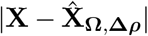. The right column Figures C and F show the contributions from spatial interactions only,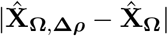.

The connection between *Mbp* and *Flt1* was interesting for a number of reasons. Myelin basic protein is the second most abundant protein, after proteolipid protein, of the myelin membrane in the central nervous system (CNS), making up approximately 30% of the total protein of myelin [30] and *Mbp* mutants lack compact myelin in the CNS [31]. Myelin surrounds nerve cell axons to insulate them and to increase the conduction of electric impulses [32]. Demyelinating diseases of the CNS, of which multiple sclerosis (MS) is the most common, are characterized by damaged myelin sheaths [33]. *Flt1* encodes a member of the vascular endothelial growth factor receptor (VEGFR) family. VEGFRs mediate diverse cellular communication signals controlling developmental processes, such as neurogenesis or gliogenesis [34]. VEGFRs recognize vascular endothelial growth factors (VEGFs), whose expression has been shown to be upregulated in both acute and chronic MS plaques [35]. Our SpaCeNet model resolves that *Flt1* levels are negatively associated with *Mbp* levels in neighboring cells. A possible interpretation is that cells expressing *Flt1* accumulate in the spatial vicinity of cells which lack *Mbp*. This would be in line with the observation that VEGF-A (which is recognized by Flt1) promotes migration of oligodendrocyte precursor cells (OPCs) in a concentration-dependent manner [36], as shown by anti-Flk-1 (not by anti-Flt-1) receptor-blocking antibody. Moreover, it was shown in the medulla oblongata of the adult mouse that OPCs contribute to focal remyelination and that VEGF signaling might be required for their proliferation [37]. It has been long known that oligodendrocytes are the myelinating cells of the CNS derived from OPCs [38], therefore our finding could be a hint that Flt1 mediates signals necessary to guide myelinating cells such as OPCs and mature oligodendrocytes to axons that lack myelin. Interestingly, in a study by Vaquie et al. [39], injured axons were shown to communicate with Schwann cells to trigger the formation of actin spheres. These spheres constrict the axons, leading to their fragmentation and faster removal of debris after injury. This process was shown to be controlled by VEGFR1 activity, and can be acquired by oligodendrocytes through enforced expression of VEGFR1 [39].

It should be noted that SpaCeNet models are associative, not causal. Thus, we can not elucidate whether decreased *Mbp* levels imply increased *Flt1* in neighboring cells or if increased *Flt1* levels imply decreased *Mbp* in neighboring cells. Moreover, since only a selected set of 28 genes was spatially assessed by STARmap in a thick tissue section, and since numerous signaling steps may be involved in establishing this relationship, the underlying mechanisms remain to be fully elucidated.

The spatial association *Flt1* -*Mbp* was followed in strength by *Ctgf* -*Gja1, Ctgf* -*Pcp4*, and *Reln*-*Sst* associations. Connective tissue growth factor (*Ctgf*), also known as Cellular Communication Network Factor 2 (*Ccn2*), belongs to the CCN family. CCN proteins are a family of extracellular matrix proteins involved in intercellular signaling [40]. We obtained Δ*ρ*_*Ctgf, Gja1*_ = 0.103, with leading-order contribution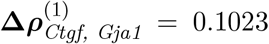. *Ctgf* has been reported to facilitate gap junction intercellular communication in chondrocytes through up-regulation of connexin 43 (*Gja1*) expression [41]. Purkinje Cell Protein 4 (Pcp4) regulates calmodulin activity and might contribute to neuronal differentiation through the activation of calmodulin-dependent kinase signaling pathways [42]. The *Ctgf* -*Pcp4* potential was estimated with Δ*ρ*_*Ctgf, Pcp4*_ = 0.083 and 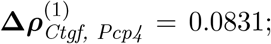; we are not aware of a mechanism that might explain the spatial association observed between *Ctgf* and *Pcp4*. This association was followed by *Reln* (Reelin) -_*Reln, Sst*_ *Sst* (Somatostatin) with parameters Δ*ρ*_*Reln, Sst*_ = 0.083 and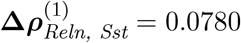. The gene *Reln* encodes a secreted extracellular matrix protein that might play a role in cell-cell interactions critical for cell positioning. Somatostatin affects transmission rates of neurons in the CNS and cell proliferation. GABAeric cortical interneurons can be delineated to 95% by the markers Pv, Sst (co-expressed with Reelin), Reelin (without Sst), and Vip [43]. The medial ganglionic eminence gives rise to the population of interneurons that co-express Reelin and Somatostatin, while the caudal ganglionic eminence gives rise to interneurons that express Somatostatin but lack Reelin [43]. The observed spatial *Reln*-*Sst* association could be a hint that the distribution of these two populations is spatially well organized and not random.

SpaCeNet assumes multivariate normality. To further ensure that the derived results are robust and to explore the extent to which they are biased by data preprocessing, we repeated the former analysis but used log-transformed data instead. Log-transformed data are believed to make data more normally distributed, but are also prone to inflating the variance for lowly expressed genes [44]. Thus, it is not *a priori* clear whether SpaCeNet can better deal with data on a natural scale or on log scale. Respective top edges obtained by SpaCeNet are summarized in Suppl. Table S5, illustrating that the top hits are still recovered with confidence, although the ordering slightly changed; among the top genes, the pairs *Mbp-Flt1, Ctgf-Gja1*, and *Ctgf-Pcp4* were confirmed. The pair *Sst-Pvalb* newly appeared among the top hits, which might be attributed to spatial delineation of Parvalbumin-positive (*Pvalb+*) and Somatostatin-positive (*Sst+*) cells.

As a final consistency check, we explored the potential role of redundant variables and how they potentially compromise model inference. For this purpose, we replaced *Mbp* in the data matrix by two noisy copies of itself and repeated our analysis with results shown in Suppl. Table S5. We observed that the edges to *Mbp* are still robustly established, but edge strength was lowered by a factor of approximately two. This is in line with recent findings in mixed graphical models [45].

#### Comparative analysis of spatial associations using pair-wise and partial spatial correlations

For comparison and illustration purposes, we performed the ordinary pair-wise spatial correlation analysis outlined above on the mouse visual cortex data, selecting *K* = 5 in line with [13]. The largest absolute spatial correlation value between different genes was observed for the pair *Mbp*-*Flt1* with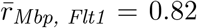, which was also the top hit in the SpaCeNet analysis. However, we observed the opposite sign for the association; SpaCeNet suggests a negative, while this analysis suggests a positive association. To further resolve this contradictory result, we compared the estimated pair-wise spatial correlation matrix with an ordinary correlation matrix derived across all single-cell profiles (see Suppl. Figures S10A and S11A), respectively. Comparison of both matrices reveals a strong coincidence with differences distributed as shown in Suppl. Figures S10 and Suppl. Figure S11.

Given the high similarity between both, we hypothesized the presence of strong auto-correlations between neighboring cells, meaning that high spatial correlations are just the consequence of similar expression patterns in adjacent cells. For the pair *Mbp*-*Flt1*, the ordinary correlation analysis revealed a correlation coefficient of 0.87, supporting this hypothesis for this specific case.

To explore if such spatial auto-correlations deteriorate estimates of spatial associations, we applied additionally the outlined spatial partial correlation measure with *K* = 5. Table 2 compares the estimated spatial partial associations with the results of SpaCeNet for the outlined top pairs. Both approaches now suggest a negative association for all four pairs. Moreover, the relative association strength is very similar with *Mbp*-*Flt1* being the strongest association, and *Ctgf* -*Pcp4* and *Reln*-*Sst* having similar association strengths. The pair *Ctgf* -*Gja1* shows a relatively weak association in contrast to the SpaCeNet analysis. One should note, however, that previous analysis was based on a nearest neighbor estimate of gene expression, ignoring potential complex radial dependencies, and did not explore the full capacity of PGMs to disentangle direct from indirect relationships by considering all molecular variables simultaneously, which might explain the observed differences. Interestingly, ranking from highest to lowest absolute spatial partial correlations also yielded the pair *Mbp*-*Flt1* as top hit, suggesting that disentangling the intra- from inter-cellular correlations might be one of the main ingredients necessary to identify the most interesting spatial associations.

**Table 2:** Estimated spatial partial correlations versus the association strength determined by SpaCeNet for the four top-ranked associations discussed in the main text. Here, *jk*^*′*^ corresponds to spatial partial correlations between variable *j* and *k*^*′*^ conditioning on *j*^*′*^ and *k*. An analoguous definition holds for *j*^*′*^*k* and the respective average is given by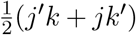. SpaCeNet’s leading order association strength is given by **Δ*ρ***^(1)^ with positive signs corresponding to negative associations in line with the definition of the precision matrix of ordinary GGMs.

| | $jk'$ | $j'k$ | $\frac{1}{2}(j'k + jk')$ | $\Delta \rho^{(1)}$ |
| --- | --- | --- | --- | --- |
| Flt1 - Mbp | -0.560 | -0.557 | -0.558 | 0.327 |
| Ctgf - Gja1 | -0.118 | -0.119 | -0.119 | 0.102 |
| Ctgf - Pcp4 | -0.227 | -0.234 | -0.231 | 0.083 |
| Reln - Sst | -0.174 | -0.205 | -0.190 | 0.078 |

### 2.3 SpaCeNet infers spatial gene-gene association patterns in the ***Drosophila*** blastoderm from low-resolution spatial omics

The previous analysis used data generated by STARmap [11], an in-situ sequencing technology for dense measurements at single-cell resolution, which, however, is limited in throughput.

In contrast, omics readouts together with spatial information (the transcriptome in a given spatially defined region of a specimen) have become more and more available. Here, we illustrate that SpaCeNet augments their analysis, even though the data do not resolve single cells.

The Berkeley Drosophila Transcription Network Project used a registration technique that uses image-based data from hundreds of *Drosophila* blastoderm embryos, each co-stained for a reference gene and one gene out of a preselected gene set, to generate a virtual *Drosophila* embryo [46]. We retrieved these virtual embryo data consisting of 84 genes whose expression levels were measured at 3039 embryonic locations from http://bimsbstatic.mdc-berlin.de/rajewsky/DVEX/. We then used SpaCeNet to estimate the intracellular network (the precision matrix **Ω**) and the network of spatial molecular interactions (**Δ*ρ***^*(·)*^) for *L* = 3. We performed a hyper-parameter grid search, where we trained the model on 70% of the data and validated it on the remaining 30% (Suppl. Figure S12 and S13). The best set of hyper-parameters was selected based on the highest validation pseudo-log-likelihood and was used to fit a final model on the full data (Suppl. File 2). The intracellular network **Ω** is a full matrix and so there is a rich dependency structure among variables not related to their spatial context. The spatial-interaction network, **Δ*ρ***^*(·)*^, in contrast, is sparse with 238 out of 3570 possible interactions (Suppl. Figure S14). Note that the spatial molecular interactions improved the goodness-of-fit on validation data and thus also the generalizability of the model, which highlights the need to include spatial interactions.

Figure 4 contrasts the expression of the gene pairs *sna*-*twi, ems*-*noc* and *Dfd*-*lok*, that showed the highest spatial association in the SpaCeNet analysis with Δ*ρ*_*twi, sna*_ = 0.0368, Δ*ρ*_*ems, noc*_ = 0.0306, and Δ*ρ*_*Dfd, lok*_ = 0.0211. We observed *twi* and *sna* to be active in the same spatial regions, which is consistent with both the positive spatial association suggested by SpaCeNet in leading order 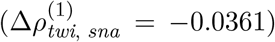 and with the joint activation of *twi* and *sna* in the differentiation of the *Drosophila* mesoderm in localized (ventral) regions of early embryos [47]. In contrast, the genes *ems* and *noc* (for which SpaCeNet estimated a negative spatial association at leading order 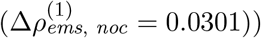 are active in adjacent but different areas of the *Drosophila* embryo. A similar observation can be made for the genes *Dfd* and *lok*, for which SpaCeNet also estimated a negative leading-order association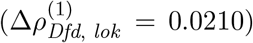. To further explore the robustness of former results with respect to pre-processing biases, we re-performed our analysis but additionally applied a log-transformation (Suppl. Table S6). Again, these analyses support the robustness of SpaCeNet as almost all leading edges coincide between both approaches.

**Figure 4:**
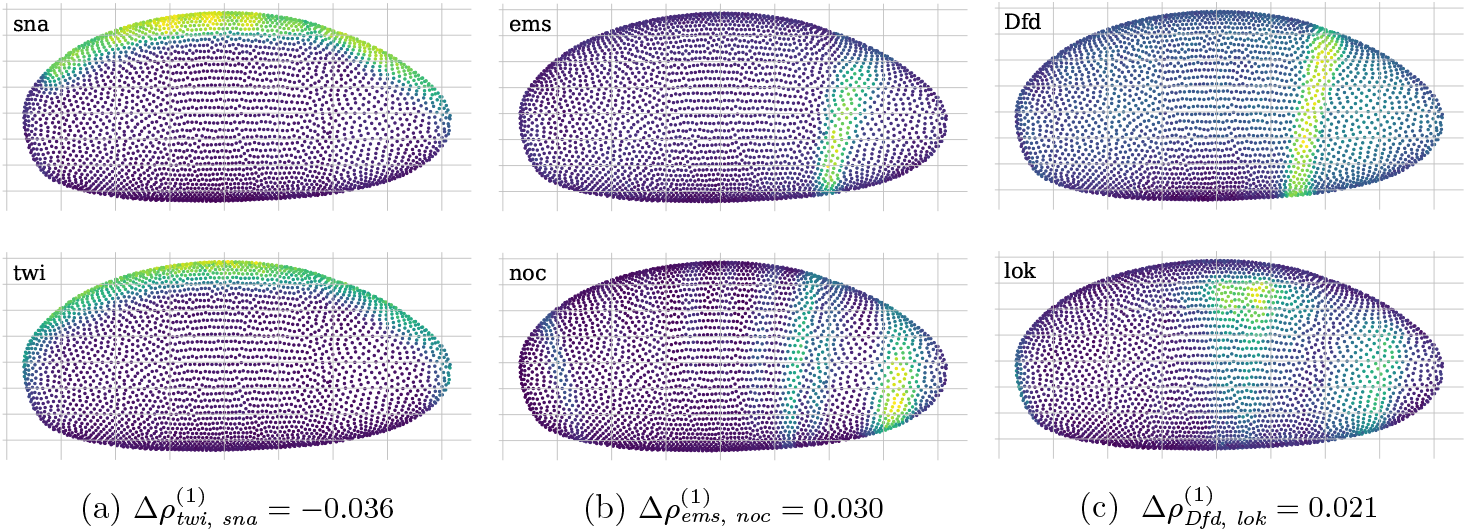
Visualization of genes with strong spatial associations in the *Drosophila* blastoderm as identified by SpaCeNet. Figures show gene expression levels of the top ranked gene pairs as identified by SpaCeNet with *sna* and *twi* in Figures A and B, *ems* and *noc* in Figures C and D, and *Dfd* and *lok* in Figures E and F, respectively. The genes *sna* and *twi* (Figures A and B) are expressed in the same areas in concordance with 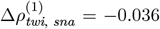 as inferred by SpaCeNet. Genes *ems* and *noc* (Figures C and D) are expressed in adjacent but different areas, which is consistent with the inferred SpaCeNet association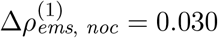. This also holds true for the gene pair *Dfd* and *lok* (Figures E and F), which are also expressed in adjacent but different areas, consistent with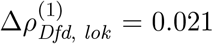. For all illustrations, the three-dimensional spatial coordinates were projected into a two-dimensional plane using a principal component analysis.

#### SpaCeNet as an inferential tool to predict spatial gene expression in the ***Drosophila*** blastoderm

SpaCeNet infers a joint density function describing spatially distributed, potentially high-dimensional molecular features. Thus, it can be also utilized as an inferential tool, predicting gene expression of a cell given its cellular context. For illustration purposes, consider the gene *Kr* that encodes the Krüppel protein, a transcriptional repressor expressed in the center of the embryo during the cellular blastoderm stage [48]. First, we predicted *Kr* expression in a leave-one-position-out approach that provides levels of *Kr* based on a cell’s environment. The ground truth is shown in Suppl. Figure S15A and the corresponding predictions in Suppl. Figure S15B. We found that an accordingly trained SpaCeNet model can predict each cell’s individual expression, given its location in space (mean squared error (MSE) = 4.36 *·* 10^*−*3^). However, slight deviations were observed in the highlighted region (red arrow). Next, we tested whether this discrepancy could be resolved by specifying *a priori* the expression levels of the remaining genes at the position of interest (Suppl. Figure S15C). This reduced the MSE to 2.88 *·* 10^*−*3^, and the highlighted region better agrees with the ground truth, as can be seen by comparing Suppl. Figure S15A to C.

#### SpaCeNet analysis of high-throughput spatial transcriptomics

Finally, we illustrate that SpaCeNet can deal with high-throughput data, capturing hundreds or even thousands of variables and measurements for tens of thousands of cells. In [49], the mouse organogenesis spatiotemporal transcriptomic atlas (MOSTA) was generated, which provides gene-expression maps with single-cell resolution of mouse organogenesis. We downloaded pre-processed data of the coronal hemibrain section, capturing in total measurements of 56,731 individual cells together with spatial locations^1^. We further filtered the gene space to genes which are at least expressed in 30% (Mosta A) and 10% of cells (Mosta B), yielding 315 and 1741 genes, respectively. We then applied SpaCeNet to each of these datasets (using *L* = 1 for computational efficiency), and performed a grid search for model selection. The best model was selected based on smallest Akaike Information Criterion (AIC). The top spatial associations are summarized in Suppl. Table S7 for. The total computation time for the final model was approximately 30 and 150 minutes, respectively, computed on a single A100 GPU, highlighting that SpaCeNet is computationally highly efficient. We want to highlight that model findings might be affected by low sequencing depths, challenging SpaCeNet’s model assumptions. However, despite this limitation given by the data, we want to discuss the findings. For both datasets Mosta A and B, the top hits predominantly showed a positive spatial association (Δ*ρ*_*ij*_ *<* 0). For instance, among the top 5 hits in both analysis, we observed the gene pairs *Mt1-Mt2* and *Mobp-Mbp. Mt1* and *Mt2* are both melatonin receptors. Interestingly, analyses in adult rat brain showed differences in the distribution of MT1 and MT2 proteins, and the labeling often appeared complementary in regions displaying both receptors [50]. The positive spatial association observed by SpaCeNet would be in line with this finding; a high expression of MT1 in a cell implies a high expression of MT2 in its neighbors and vice versa. Myelin-associated oligodendrocytic basic protein (Mobp) is a myelin constituent exlusively expressed by oligodendrocytes [51] and shares physiochemical and biological properties with Mbp. Both were expressed in similar regions (Suppl. Figure S17), but were shown to differ with respect to localization and expression timing [51], with Mobp occurring significantly later than Mbp at the late stages of myelination. Finally, considering negative spatial associations, we detected a strong connection between *Apoe* and *Ptgds*. Prostaglandin-H2 D-isomerase is an enzyme encoded by the *Ptgds* gene. Prostaglandins are key players in neuroinflammatory and neurodegenerative diseases [52]. Apolipoprotein E (*Apoe*) influences various biological processes, and both the *Apoe* gene and its encoded proteins hold promise as targets for therapies against Alzheimer disease (AD) [53]. In summary, even in scenarios where SpaCeNet’s model assumptions are challenged, it suggests promising findings for further analyses.

## 3 Discussion

SpaCeNet is a network inference method that determines intracellular correlation networks and cell-cell associations from spatial molecular data. SpaCeNet is based on probabilistic graphical modeling and extends the concept of conditional independence (CI) by spatial information through estimating spatial conditional independencies (SCI). These intracellular and intercellular SCI relationships encode information about how molecular variables affect each other across space. We verified SpaCeNet in comprehensive simulation studies and demonstrated the information that SpaCeNet can extract from spatial transcriptomics data in two example data sets: an expression map of the mouse visual cortex and a virtual RNA map of the *Drosophila* blastoderm. The analysis of the mouse visual cortex allowed us to generate hypotheses about the spatial organization of cell populations, such as *Flt1* -mediated signals which could be involved in the recruitment of myelinating cells towards axons which lack myelin basic protein (*Mbp*). The analysis of the *Drosophila* blastoderm showed that SpaCeNet can also yield insights if data do not have cellular resolution. In the latter case, SpaCeNet resolved spatial association patterns between molecular variables, as observed between the gene-pairs *twi*-*sna* and *ems*-*noc*.

Modern spatial transcriptomics techniques are capable of measuring RNA transcript levels with single-cell resolution in a three-dimensional space. Many molecular processes that take place within and between cells, such as translation to protein, possible post-translational modifications, and subsequent signaling cascades comprising secretion of molecules and their recognition by other cells, are not directly measured by these technologies. Although laboratory methods under development, such as single-cell proteomics and metabolomics [54, 55] or single-cell epigenetic or chromatin confirmation measurements may one day bridge this gap, at present one can only infer such associations with appropriate analytical methods. SpaCeNet is an important first step in inferring complex, multi-omic, intracellular and intercellular association networks that takes advantage of high-dimensional omics data and spatial information. The present implementation of SpaCeNet uses single-cell spatial transcriptomics data and so can only infer associations between RNA expression levels within and, more importantly, between different cells – without knowledge about any potentially complex intermediate processes. SpaCeNet is based on a robust Gaussian graphical model framework based on partial correlations, which can scale to the increasingly large and complex spatial multi-omics data sets that new laboratory technologies provide. Importantly, hidden variables can prohibit the inference of direct relationships. If they remain hidden, we are not able to infer their role in mediating dependencies, potentially leading to erroneous direct associations. However, with new technologies capturing more and more molecular variables, methods such as SpaCeNet might be key to identify the direct statistical dependencies in favor of those mediated by other variables. In this context, it is also worth mentioning that SpaCeNet’s current implementation is limited to modeling continuous variables. While many omics measurements such as gene expression levels can be reasonably modeled as continuous, this is not the case for all genomic readouts, and variables such as mutation status and chromatin accessibility might be better described as categorical variables. SpaCeNet could be easily adapted to include those using mixed graphical modeling suggested by [56]. As is generally the case for correlation-based methods, SpaCeNet does not identify causal interactions, although lower bounds on causal effects can be derived from observational data [29], and techniques such as directed acyclic graphs and respective equivalence classes may be adapted to SCI relationships, which we intend to explore in the future.

Nevertheless, SpaCeNet represents an important step forward in the analysis of spatial expression data, allowing us to move from a simple atlas of expression values and cell types to models that capture complex patterns of interactions that allow tissues to function and guide cellular growth, development, and disease processes. As new experimental techniques deliver larger and more complex multi-omics data combined with higher resolution information on the location of individual cells, techniques like SpaCeNet will become increasingly important for integrating spatial and biological contexts.

## 4 Materials and Methods

### 4.1 Data structure

SpaCeNet is designed to model continuous, potentially high-dimensional molecular variables (e.g., gene-expression levels) which are measured together with spatial information. Formally, denote the molecular variables as *X*_*i*_ with *i* = 1, …, *p* and let **X** = [**x**^1^, …, **x**^*n*^]^*⊤*^ *∈* ℝ^*n×p*^ be a data matrix with *n* measurements (molecular profiles) in its rows. Then each of these measurements **x**^*a*^ is annotated with a position *r*_*a*_ of cell *a* in space. Throughout the section, we identify the **x**^*a*^ as molecular profiles of individual cells and *r*_*a*_ with the respective point-position in space.

### 4.2 Concept

SpaCeNet considers the observed spatially distributed molecular profiles of single cells within a statistical model which decomposes the observed gene expression into two components (see Figure 1). The first contribution accounts for the statistical diversity of isolated cells, meaning that we assume that the molecular profiles of individual cells follow a *p*-dimensional multivariate normal distribution, *X ∼ N* (*μ*, Σ) with mean profiles *μ* and covariance Σ (see Figure 1A). Note that this is a reasonable assumption if data are approximate normal or if they are transformed accordingly (e.g., via non-paranormal transformations) [57, 58]. The second contribution accounts for the fact that cells are not isolated but observed in a complex environment where cells can interact with each other (see Figure 1B and C). This effect is encoded as shifts of the distribution’s mean profiles *μ* and the amount by which the individual profiles are shifted depends on the molecular phenotypes of the surrounding cells and their distances, as schematically illustrated for two interacting cells in Figure 1B and for multiple cells in 1C. The functional dependence of these shifts on the neighbor’s gene expression and distance is not a priori clear. Key to SpaCeNet is a parameterization which accounts for diverse distance dependencies and which can be straightforwardly interpreted in terms of conditional independence relationships, as outlined in the following.

### 4.3 Spatial conditional independence

Estimates of Conditional Independence (CI) relationships are key to the inference of molecular networks, as they allow the disentanglement of direct and indirect statistical relationships and as such reduce the number of false positive associations [25]. If the spatial context is ignored, CI between variables *X*_*i*_ and *X*_*j*_ given all remaining variables can be expressed as *X*_*i*_ *⊥ X*_*j*_|*{*rest*}*, where “rest” is the set of all variables in *X* except *X*_*i*_ and *X*_*j*_. We extend this language to intracellular and intercellular spatial conditional independence (SCI) relationships:

- *Intracellular SCI* relations between variables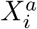 and 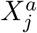measured within one cell *a* are expressed as 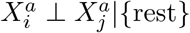with the term “rest” referring to all other variables of cell *a* and to all variables of all other cells.
- *Intercellular SCI* relations between variables 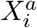 and 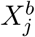 measured in different cells *a* and *b*, conditioned on all remaining variables, are denoted as 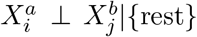with *a≠ b*, where “rest” refers to all variables of cell *a* except 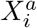, all variables of cell *b* except 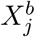 and all variables of all remaining cells.

Thus, such as CI is designed to distinguish direct from indirect relationships between molecular variables, SCI relationships are designed to do so for the intra- and intercellular relationships between molecular variables.

### 4.4 Full probability density

Gaussian Graphical Models (GGMs) are PGMs which assume multivariate normally distributed data. The model parameters of GGMs are collected in the precision matrix **Ω** = (*ω*_*ij*_), which directly parameterizes CI relationships; *ω*_*ij*_ = 0 if and only if there is CI between variables *i* and *j* given all other variables. In the following, we will develop a joint probability density which directly encodes SCI relationships. Specifically, we assume that data follow an *np*-dimensional multivariate normal distribution with precision matrix **Λ**, where the entries in **Λ** capture both intracellular and intercellular SCI relations. The *full probability density* of SpaCeNet is

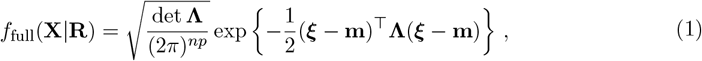

where we stack the individual cells’ profiles **x**^*a*^ *∈* ℝ^*p*^, *a* = 1, …, *n*, vertically in ***ξ*** = vec(**X**^*⊤*^) and use a global, location-agnostic mean vector ***μ*** for all **x**^*a*^ such that **m** = **1**_*n*_ *⊗* ***μ*** = (*μ*_1_, …, *μ*_*p*_, *μ*_1_, …, *μ*_*p*_, …)^*⊤*^. All pairwise cell-cell distances are collected in a matrix **R** = (*r*_*ab*_) *∈* ℝ^*n×n*^, where *r*_*ab*_ = *r*_*ba*_ denotes the Euclidean distance between cells *a* and *b*. The space-agnostic conditional independence (CI) relations for gene expression levels of a single cell *a* can be recovered from Eq. (1) by marginalizing over all variables of all other cells *b ≠ a*.

Next, we decompose the precision matrix **Λ** *∈* ℝ^*np×np*^ into

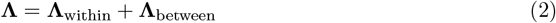

with a matrix for intracellular (within-cell) associations

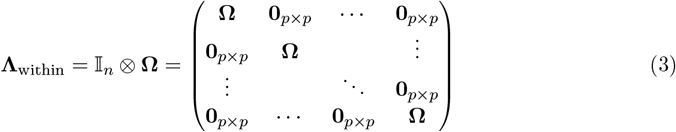

and a matrix for intercellular (between-cell) associations

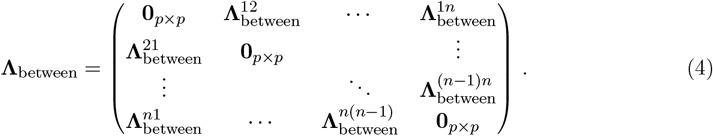

Due to the parametrization of **Λ**_within_ in Eq. (3), the same precision matrix **Ω** = (*ω*_*ij*_) *∈* ℝ ^*p×p*^ parametrizes the within-cell associations for all cells, where *ω*_*ij*_ = 0 corresponds to intracellular SCI between *X*_*i*_ and *X*_*j*_. The conventional (space-agnostic) GGM is recovered for **Λ**_between_ = **0**_*np×np*_.

We further assume that the intercellular association of gene *i* in any cell *a* with gene *j* in any other cell *b* is described by some function of the cells’ Euclidean distance *r*_*ab*_. We denote this radial cell-cell interaction potential by *ρ*_*ij*_(*r*_*ab*_) and write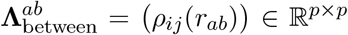, which we require to be symmetric with respect to both *i ↔ j* and *a ↔ b*. The (*np × np*)-dimensional precision matrix **Λ** is required to be positive definite for *f*_full_ to be a valid probability density.

### 4.5 Cell-cell interaction potentials

From the definition of the probability density function, Eq. (1), we see that

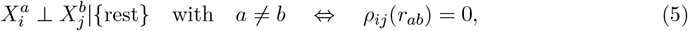

where the set “rest” refers to all variables of cell *a* except 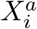, all variables of cell *b* except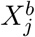, and all variables of all other cells. Thus, all intercellular SCI relations are encoded in the set of *p*(*p* + 1)*/*2 independent functions *ρ*_*ij*_(*r*) = *ρ*_*ji*_(*r*). To ensure that cells that are infinitely separated do not interact, we require *ρ*_*ij*_(*r*) = 0 for *r → ∞*. To approximate the potential *ρ*_*ij*_(*r*), we use a power-series in 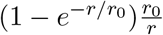,

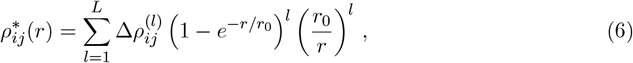

where *l* is the order in the series expansion and 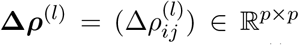the corresponding coefficient matrix. The latter matrices are required to be symmetric, 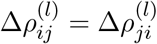. This leads to an approximation for the intercellular precision matrix **Λ**_between_ given by

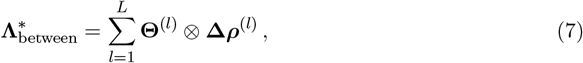

with

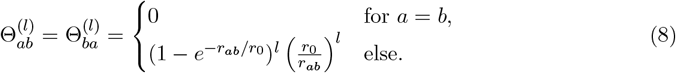

The coefficients 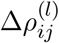are model parameters which are estimated using a regularized pseudo-log-likelihood, as outlined below. The expansion (6) naturally fulfills 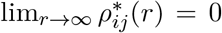 and 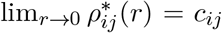 with constants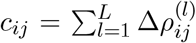. Note, an expansion in 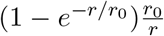 has the advantage that terms do not diverge for *r →* 0, which is in contrast to an expansion in 1*/r*. Thus, the factor (1*−e*^*−r/r*0^) smooths the divergence and the amount of smoothing is determined by the additional parameter *r*_0_.

### 4.6 Pseudo-log-likelihood

The precision matrix **Λ** is of size *np × np*, which makes a naive maximum-likelihood-based estimate (using, for example, a gradient descent) intractable for reasonably large *p* and *n*. We address this issue by using a pseudo-log-likelihood approach [59, 56], which is a computationally efficient and consistent estimator formed by products of all the conditional distributions. Let **X**^**\a**^ denote all gene expression levels in all cells except cell *a*, and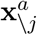 denote all gene expression levels in cell *a* except gene *j*. We consider the conditional density

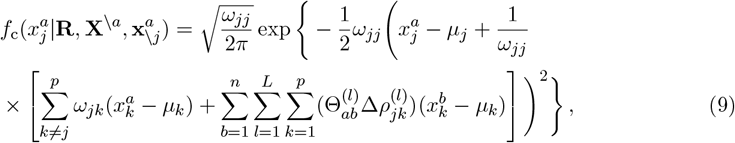

obtained from the full density given in Eq. (1). This yields the pseudo-log-likelihood

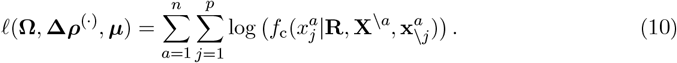

### 4.7 Regularization

Parameter regularization has been repeatedly shown to improve the inference of GGMs using approaches such as node-wise lasso regression [60], the graphical lasso [61], or covariance shrinkage [62]. This is particularly the case if the number of variables exceeds or is of the same order of magnitude as the number of measurements [25]. For *S* independent measurements (spatial transcriptomics slides), the full, regularized pseudo-log-likelihood-based optimization problem of SpaCeNet is given by

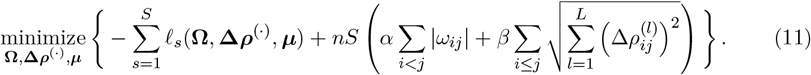

Eq. (11) penalizes the off-diagonal elements of the intracellular precision matrix **Ω** via *L*_1_ regularization [63]. The intercellular interactions are regularized via group-lasso terms [64], where the group contains the coefficients of the spatial interactions at different orders *l* = 1, …, *L* of the series expansion. This regularization has the advantage that sparseness is induced simultaneously across all orders in the expansion, so we induce sparseness in the potentials *ρ*_*ij*_ and not just in the different orders of its expansion. The hyper-parameters *α* and *β* calibrate the regularization strength of the intracellular and intercellular associations, respectively. In our analysis, we standardized the expression data prior to model learning to ensure comparable penalization of the variables.

### 4.8 Implementation

Eq. (11) is a convex optimization problem with lasso and group-lasso regularization terms, which can be efficiently solved via proximal gradient descent [65]. SpaCeNet uses a proximal gradient descent with Nesterov acceleration [66, 67]. The optimization terminates when the loss in Eq. (11) improves less than a user-specified threshold. The computational requirements for training a SpaCeNet depend on the number of considered cells and genes. For example, training SpaCeNet on a Nvidia A100 GPU takes around one minute for the Drosophila data (3.039 cells and 84 genes), less than two minutes on the StarMap data (33.598 cells and 28 genes), around 30 minutes on the Mosta data with more than 30% non-zero entries (56.731 cells and 315 genes) and around 150 minutes on the Mosta data with more than 10% non-zero entries (56.731 cells and 1741 genes).

## 5 Supplementary Data

Supplementary Data are available online.

## Footnotes

1 https://ftp.cngb.org/pub/SciRAID/stomics/STDS0000058/Cell_bin_matrix/Mouse_brain_Adult_GEM_CellBin.tsv.gz

