## Supplementary material for "SpaCeNet: Spatial Cellular Networks from omics data"

### Supplementary Material to
### "SpaCeNet: Spatial Cellular Networks from omics data"

#### 1 Simulation studies

**Data simulation** Data were simulated from the full probability density (main article, Eq. (1)) with potentials $\rho_{ij}(r_{ab}) = \Delta\rho_{ij}\exp(-\phi_{ij} r_{ab})$, using the following procedure:

1. Initialize empty $p \times p$ matrices for $\mathbf{\Omega}$ and $\mathbf{\Delta\rho}$.
2. Randomly draw a set of symmetric edges (we chose 5% of all possible edges) for $\mathbf{\Delta\rho}$ from Unif$(-1, 1)$ and draw corresponding range parameters $\phi_{ij} = \phi_{ji}$ for given positive $\phi_{min}$ and $\phi_{max}$ from Unif$(\phi_{min}, \phi_{max})$.
3. Randomly draw a set of symmetric edges (we chose 10% of all possible edges) for $\mathbf{\Omega}$ from Unif$(-1, 1)$.
4. Draw $s = 1, \ldots, S$ samples of spatial coordinates, uniformly sampled from 3d space with an average density of 100 cells per volume unit, each with $n$ cells and calculate the pairwise distances $\mathbf{R}^{(s)}$ for each sample.
5. Based on $\mathbf{\Omega}$, $\mathbf{\Delta\rho}$, $\phi_{ij}$ and $\mathbf{R}^{(s)}$ construct $\mathbf{\Lambda}^{(s)}$ for all $s$.
6. Calculate the row-wise sum of the absolute values of all $\mathbf{\Lambda}^{(s)}$. For each of the $p$ variables, select the maximum value from the corresponding $n \cdot S$ sums, add a small constant (we chose $10^{-7}$) and fill the respective diagonal element of $\mathbf{\Omega}$ with it to ensure that $\Lambda$ is positive definite.
7. Update all $\mathbf{\Lambda}^{(s)}$, sample $\boldsymbol{\xi}^{(s)}$ from $\mathbf{\Lambda}^{(s)}$ by means of a Cholesky decomposition and add a random $\boldsymbol{\mu}$ if desired.

With this setup, we simulated 24 scenarios with different parameter settings for $n$, $S$ and $\phi_{ij}$ with 20 independently seeded replicates for each. All simulations used $p = 20$ variables. Note that the data are simulated from the full joint probability Eq. (1) of the main manuscript with a precision matrix of size $pn \times pn$, making data simulation computationally expensive for large $p$ and $n$.

In our studies, $\phi_{ij}$ was chosen between 5 and 20. The rationale behind this choice is motivated as follows. Given a unit density of $\eta$, $k$ cells on average occupy a volume $V_k = k/\eta$, which corresponds to a sphere of radius $r_k = (3k/(4\pi\eta))^{1/3}$. We used $r_k$ as an estimate for the average distance between a cell and its $k$ nearest neighbors. For $\eta = 100$, this yields $r_1 \approx 0.13$, $r_8 = 2r_1 \approx 0.27$, and $r_{27} = 3r_1 \approx 0.40$. The range of the potentials $\exp(-\phi_{ij} r)$ can be quantified by $1/\phi_{ij}$. With the average nearest-neighbor distance $r_1$ as reference point, $\phi_{ij} = 5$ therefore corresponds to a long-range and $\phi_{ij} = 20$ to a short-range potential.

**SpaCeNet model selection** We set the expansion order to $L = 3$ and chose $r_0$ to equal the minimal observed distance between two cells. Note that there are reasonable alternative choices for $r_0$, e.g., the average nearest-neighbor distance provides a length scale that might be more appropriate for larger numbers of cells $n$. We then performed a grid search on all data sets with 4 values for $\alpha \in [10^{-5}, 10^{-3}, 10^{-1}, 10]$ and $\beta \in [10^{-5}, 10^{-3}, 10^{-1}, 10]$ each. The grid was then successively refined 6 times such that about 100 different hyper-parameter combinations were evaluated in total. The best set of hyper-parameters was chosen based on the maximum pseudo-log-likelihood of test data. To this end, the full data set was split 70:30 into a training and a test set. If more than one sample was available ($S > 1$), the split was performed between different samples and otherwise ($S = 1$) all observations in 30 % of the spatial volume were used for testing. We initialized the optimization with a step size of $10^{-6}$. If overflows were encountered, we successively reduced the step size by a factor of 10. The convergence threshold for the proximal gradient algorithm was set to $10^{-5}$ and training was terminated after a maximum of 3,088 optimization steps. The AUROC and AUPRC based evaluation methods rely on a threshold-based classification of estimated parameters into positives and negatives. When considering the spatial association parameters we classify an association between two variables $i$ and $j$ to be positive if $|\Delta\rho_{ij}^{(l)}|$ is greater than the threshold for at least one considered order $l$ in the expansion.

**Reconstruction of interaction potentials** Data for the reconstruction of the interaction potentials (main article Figure 2) was generated in line with previous procedure, but using $p = 5$ and only a single spatial edge connecting two of the variables with the potentials given in the caption of main article Figure 2a to d, respectively. We simulated data for $n = 10$ and $S = 1000$, and the potentials were fitted with $r_0 = 0.1$.

### 2 Estimates from conditional densities

By construction, we assume a spatially constant mean vector $\hat{\mathbf{x}} = \boldsymbol{\mu}$ for all cells in the full density distribution of SpaCeNet. Then, we directly obtain from the conditional densities that

$$\hat{x}_j^a = \mu_j - \frac{1}{\omega_{jj}} \left[ \sum_{k \neq j}^{p} \omega_{jk}(x_k^a - \mu_k) + \sum_{b=1}^{n} \sum_{l=1}^{L} \sum_{k=1}^{p} (\Theta_{ab}^{(l)} \Delta\rho_{jk}^{(l)})(x_k^b - \mu_k) \right] , \tag{1}$$

where the mean of the normal distribution is shifted. Eq. 1 can be used as an estimate for the variable $x_j^a$ provided all other variables are known.

This method was used to calculate the residuals for the mouse visual cortex data (main article Fig. 4), where $\boldsymbol{\Delta\rho}^{(\cdot)}$ was either set to 0 or the estimated parameters. In this case, information $\mathbf{x}_{\setminus j}^a$ is required, i.e., all other variables of cell $a$ have to be known already.

Similarly, the joint conditional density for all variables of one cell $a$ is given by

$$\begin{aligned} f(\mathbf{x}^a|\mathbf{R}, \mathbf{X}^{\setminus a}) &= \frac{\sqrt{|\boldsymbol{\Omega}|}}{(2\pi)^{p/2}} \exp\left\{ -\frac{1}{2}(\mathbf{x}^a - \boldsymbol{\mu} + \boldsymbol{\Omega}^{-1}\boldsymbol{\rho}^a)^\top \boldsymbol{\Omega}(\mathbf{x}^a - \boldsymbol{\mu} + \boldsymbol{\Omega}^{-1}\boldsymbol{\rho}^a) \right\} \\ &= \frac{\sqrt{|\boldsymbol{\Omega}|}}{(2\pi)^{p/2}} \exp\left\{ -\frac{1}{2}(\mathbf{x}^a - \hat{\mathbf{x}}^a)^\top \boldsymbol{\Omega}(\mathbf{x}^a - \hat{\mathbf{x}}^a) \right\} \end{aligned}$$

with

$$\hat{\mathbf{x}}^a = \boldsymbol{\mu} - \boldsymbol{\Omega}^{-1}\boldsymbol{\rho}^a \,,$$
$$\rho_j^a = \sum_{l=1}^{L}\sum_{k=1}^{p}\sum_{b=1}^{n}\Theta_{ab}^{(l)}(x_k^b - \mu_k)\Delta\rho_{jk}^{(l)} \,.$$

The mean vector $\hat{\mathbf{x}}^a$ can be used as an estimate for the variables of cell $a$, provided that all variables of all other cells are known. Note that $\boldsymbol{\rho}^a$ only depends on other cells $b \neq a$ since $\Theta_{aa}^{(l)} = 0$ for all $l$. A comparison of the results obtained with the two methods is shown in main article Fig. 6.

### 3 Tables and Figures

Table S1: SpaCeNet results of simulation studies.

| | | | | AUROC $\mathbf{\Omega}$ | | AUPRC $\mathbf{\Omega}$ | | AUROC $\mathbf{\Delta\rho}$ | | AUPRC $\mathbf{\Delta\rho}$ | |
|---|---|---|---|---|---|---|---|---|---|---|---|
| | | | | mean | s.d. | mean | s.d. | mean | s.d. | mean | s.d. |
| $\phi_{\min}$ | $\phi_{\max}$ | $n \cdot S$ | $S$ | | | | | | | | |
| 5.0 | 5.0 | 1000 | 1 | 0.89 | 0.04 | 0.85 | 0.04 | 0.55 | 0.07 | 0.28 | 0.21 |
| | | | 10 | 0.91 | 0.04 | 0.88 | 0.04 | 0.63 | 0.11 | 0.27 | 0.17 |
| | | | 100 | 0.96 | 0.02 | 0.94 | 0.02 | 0.93 | 0.07 | 0.73 | 0.12 |
| | | 10000 | 10 | 0.95 | 0.02 | 0.94 | 0.02 | 0.87 | 0.10 | 0.64 | 0.16 |
| | | | 100 | 0.97 | 0.02 | 0.96 | 0.02 | 0.94 | 0.06 | 0.83 | 0.11 |
| | | 100000 | 100 | 0.98 | 0.02 | 0.98 | 0.02 | 0.98 | 0.03 | 0.92 | 0.12 |
| | 20.0 | 1000 | 1 | 0.96 | 0.02 | 0.94 | 0.03 | 0.59 | 0.10 | 0.25 | 0.17 |
| | | | 10 | 0.97 | 0.02 | 0.95 | 0.02 | 0.62 | 0.09 | 0.26 | 0.17 |
| | | | 100 | 0.97 | 0.01 | 0.96 | 0.02 | 0.78 | 0.10 | 0.40 | 0.12 |
| | | 10000 | 10 | 0.98 | 0.02 | 0.98 | 0.02 | 0.84 | 0.13 | 0.64 | 0.15 |
| | | | 100 | 0.99 | 0.01 | 0.98 | 0.01 | 0.90 | 0.06 | 0.75 | 0.14 |
| | | 100000 | 100 | 0.99 | 0.01 | 0.99 | 0.01 | 0.96 | 0.06 | 0.88 | 0.13 |
| 10.0 | 10.0 | 1000 | 1 | 0.95 | 0.02 | 0.93 | 0.02 | 0.62 | 0.10 | 0.24 | 0.15 |
| | | | 10 | 0.96 | 0.02 | 0.94 | 0.03 | 0.63 | 0.09 | 0.25 | 0.16 |
| | | | 100 | 0.97 | 0.01 | 0.96 | 0.02 | 0.84 | 0.12 | 0.48 | 0.13 |
| | | 10000 | 10 | 0.98 | 0.01 | 0.98 | 0.02 | 0.88 | 0.08 | 0.72 | 0.13 |
| | | | 100 | 0.99 | 0.01 | 0.98 | 0.01 | 0.93 | 0.07 | 0.81 | 0.12 |
| | | 100000 | 100 | 0.99 | 0.01 | 0.99 | 0.01 | 0.98 | 0.02 | 0.92 | 0.11 |
| 20.0 | 20.0 | 1000 | 1 | 0.98 | 0.02 | 0.97 | 0.02 | 0.57 | 0.08 | 0.20 | 0.17 |
| | | | 10 | 0.98 | 0.02 | 0.97 | 0.02 | 0.55 | 0.06 | 0.22 | 0.20 |
| | | | 100 | 0.98 | 0.01 | 0.97 | 0.02 | 0.67 | 0.10 | 0.26 | 0.13 |
| | | 10000 | 10 | 0.99 | 0.01 | 0.99 | 0.01 | 0.76 | 0.15 | 0.49 | 0.16 |
| | | | 100 | 1.00 | 0.01 | 0.99 | 0.01 | 0.88 | 0.09 | 0.65 | 0.14 |
| | | 100000 | 100 | 1.00 | 0.01 | 1.00 | 0.01 | 0.94 | 0.07 | 0.84 | 0.14 |

Table S2: Spatial edge recovery in terms of AUPRC for the correlation baseline for a varying number of nearest neighbours considered for the spatial environment

| | | | | 1knn $\boldsymbol{\Delta\rho}$ | | 5knn $\boldsymbol{\Delta\rho}$ | | 10knn $\boldsymbol{\Delta\rho}$ | | 20knn $\boldsymbol{\Delta\rho}$ | | 50knn $\boldsymbol{\Delta\rho}$ | |
|---|---|---|---|---|---|---|---|---|---|---|---|---|---|
| | | | | mean | s.d. | mean | s.d. | mean | s.d. | mean | s.d. | mean | s.d. |
| $\phi_{\min}$ | $\phi_{\max}$ | $n \cdot S$ | $S$ | | | | | | | | | | |
| 5.0 | 5.0 | 1000 | 1 | 0.07 | 0.05 | 0.08 | 0.06 | 0.08 | 0.05 | 0.07 | 0.03 | 0.07 | 0.04 |
| | | | 10 | 0.06 | 0.03 | 0.07 | 0.05 | 0.08 | 0.05 | 0.07 | 0.04 | 0.07 | 0.04 |
| | | | 100 | 0.18 | 0.10 | 0.33 | 0.14 | 0.33 | 0.14 | 0.33 | 0.14 | 0.33 | 0.14 |
| | | 10000 | 10 | 0.10 | 0.07 | 0.16 | 0.09 | 0.24 | 0.12 | 0.23 | 0.14 | 0.25 | 0.12 |
| | | | 100 | 0.22 | 0.12 | 0.48 | 0.17 | 0.55 | 0.20 | 0.57 | 0.20 | 0.43 | 0.15 |
| | | 100000 | 100 | 0.62 | 0.16 | 0.81 | 0.10 | 0.85 | 0.11 | 0.88 | 0.10 | 0.85 | 0.10 |
| | 20.0 | 1000 | 1 | 0.12 | 0.08 | 0.14 | 0.08 | 0.10 | 0.05 | 0.08 | 0.04 | 0.07 | 0.04 |
| | | | 10 | 0.07 | 0.04 | 0.08 | 0.06 | 0.09 | 0.05 | 0.07 | 0.03 | 0.06 | 0.03 |
| | | | 100 | 0.15 | 0.11 | 0.11 | 0.07 | 0.11 | 0.07 | 0.11 | 0.07 | 0.11 | 0.07 |
| | | 10000 | 10 | 0.39 | 0.19 | 0.39 | 0.15 | 0.39 | 0.19 | 0.25 | 0.16 | 0.18 | 0.11 |
| | | | 100 | 0.42 | 0.19 | 0.53 | 0.19 | 0.48 | 0.19 | 0.39 | 0.16 | 0.18 | 0.10 |
| | | 100000 | 100 | 0.84 | 0.13 | 0.84 | 0.10 | 0.80 | 0.12 | 0.76 | 0.12 | 0.64 | 0.12 |
| 10.0 | 10.0 | 1000 | 1 | 0.13 | 0.08 | 0.16 | 0.09 | 0.11 | 0.05 | 0.10 | 0.04 | 0.07 | 0.04 |
| | | | 10 | 0.07 | 0.04 | 0.09 | 0.06 | 0.09 | 0.05 | 0.07 | 0.04 | 0.06 | 0.03 |
| | | | 100 | 0.16 | 0.12 | 0.12 | 0.07 | 0.12 | 0.07 | 0.12 | 0.07 | 0.12 | 0.07 |
| | | 10000 | 10 | 0.41 | 0.19 | 0.48 | 0.16 | 0.48 | 0.20 | 0.33 | 0.17 | 0.24 | 0.13 |
| | | | 100 | 0.46 | 0.18 | 0.60 | 0.20 | 0.55 | 0.20 | 0.48 | 0.20 | 0.21 | 0.11 |
| | | 100000 | 100 | 0.88 | 0.11 | 0.89 | 0.09 | 0.89 | 0.10 | 0.88 | 0.10 | 0.81 | 0.11 |
| 20.0 | 20.0 | 1000 | 1 | 0.11 | 0.07 | 0.09 | 0.06 | 0.07 | 0.05 | 0.06 | 0.03 | 0.06 | 0.03 |
| | | | 10 | 0.07 | 0.04 | 0.07 | 0.05 | 0.07 | 0.04 | 0.06 | 0.03 | 0.06 | 0.03 |
| | | | 100 | 0.11 | 0.09 | 0.06 | 0.04 | 0.06 | 0.04 | 0.06 | 0.04 | 0.06 | 0.04 |
| | | 10000 | 10 | 0.38 | 0.18 | 0.21 | 0.13 | 0.16 | 0.10 | 0.10 | 0.11 | 0.08 | 0.05 |
| | | | 100 | 0.37 | 0.17 | 0.29 | 0.16 | 0.22 | 0.15 | 0.14 | 0.11 | 0.06 | 0.03 |
| | | 100000 | 100 | 0.81 | 0.12 | 0.78 | 0.13 | 0.69 | 0.17 | 0.55 | 0.17 | 0.31 | 0.18 |

Table S3: Spatial edge recovery in terms of AUPRC for the partial correlation baseline for a varying number of nearest neighbours considered for the spatial environment

| | | | | 1knn $\boldsymbol{\Delta\rho}$ | | 5knn $\boldsymbol{\Delta\rho}$ | | 10knn $\boldsymbol{\Delta\rho}$ | | 20knn $\boldsymbol{\Delta\rho}$ | | 50knn $\boldsymbol{\Delta\rho}$ | |
|---|---|---|---|---|---|---|---|---|---|---|---|---|---|
| | | | | mean | s.d. | mean | s.d. | mean | s.d. | mean | s.d. | mean | s.d. |
| $\phi_{\min}$ | $\phi_{\max}$ | $n \cdot S =$ | $S$ | | | | | | | | | | |
| 5.0 | 5.0 | 1000 | 1 | 0.07 | 0.05 | 0.08 | 0.05 | 0.08 | 0.05 | 0.08 | 0.05 | 0.07 | 0.04 |
| | | | 10 | 0.06 | 0.04 | 0.07 | 0.04 | 0.08 | 0.04 | 0.07 | 0.03 | 0.06 | 0.04 |
| | | | 100 | 0.18 | 0.13 | 0.32 | 0.17 | 0.32 | 0.17 | 0.32 | 0.17 | 0.32 | 0.17 |
| | | 10000 | 10 | 0.11 | 0.09 | 0.18 | 0.11 | 0.25 | 0.12 | 0.23 | 0.13 | 0.24 | 0.11 |
| | | | 100 | 0.20 | 0.11 | 0.46 | 0.17 | 0.53 | 0.20 | 0.53 | 0.18 | 0.42 | 0.15 |
| | | 100000 | 100 | 0.59 | 0.16 | 0.75 | 0.12 | 0.79 | 0.13 | 0.83 | 0.11 | 0.82 | 0.10 |
| | 20.0 | 1000 | 1 | 0.11 | 0.08 | 0.13 | 0.08 | 0.11 | 0.06 | 0.08 | 0.05 | 0.08 | 0.06 |
| | | | 10 | 0.07 | 0.05 | 0.08 | 0.05 | 0.08 | 0.05 | 0.06 | 0.03 | 0.06 | 0.04 |
| | | | 100 | 0.14 | 0.11 | 0.11 | 0.08 | 0.11 | 0.08 | 0.11 | 0.08 | 0.11 | 0.08 |
| | | 10000 | 10 | 0.40 | 0.16 | 0.37 | 0.16 | 0.37 | 0.16 | 0.24 | 0.13 | 0.16 | 0.09 |
| | | | 100 | 0.40 | 0.18 | 0.51 | 0.18 | 0.47 | 0.18 | 0.35 | 0.15 | 0.16 | 0.10 |
| | | 100000 | 100 | 0.78 | 0.15 | 0.77 | 0.13 | 0.75 | 0.15 | 0.73 | 0.16 | 0.64 | 0.19 |
| 10.0 | 10.0 | 1000 | 1 | 0.12 | 0.07 | 0.15 | 0.09 | 0.13 | 0.07 | 0.10 | 0.06 | 0.07 | 0.04 |
| | | | 10 | 0.07 | 0.05 | 0.09 | 0.06 | 0.09 | 0.05 | 0.07 | 0.04 | 0.06 | 0.04 |
| | | | 100 | 0.15 | 0.13 | 0.13 | 0.09 | 0.13 | 0.09 | 0.13 | 0.09 | 0.13 | 0.09 |
| | | 10000 | 10 | 0.41 | 0.16 | 0.46 | 0.16 | 0.46 | 0.16 | 0.33 | 0.15 | 0.21 | 0.11 |
| | | | 100 | 0.43 | 0.17 | 0.57 | 0.18 | 0.54 | 0.20 | 0.45 | 0.18 | 0.19 | 0.12 |
| | | 100000 | 100 | 0.81 | 0.13 | 0.82 | 0.11 | 0.82 | 0.12 | 0.83 | 0.11 | 0.79 | 0.11 |
| 20.0 | 20.0 | 1000 | 1 | 0.11 | 0.08 | 0.08 | 0.05 | 0.06 | 0.04 | 0.07 | 0.05 | 0.07 | 0.04 |
| | | | 10 | 0.07 | 0.05 | 0.06 | 0.04 | 0.06 | 0.03 | 0.06 | 0.03 | 0.06 | 0.04 |
| | | | 100 | 0.11 | 0.10 | 0.05 | 0.03 | 0.05 | 0.03 | 0.05 | 0.03 | 0.05 | 0.03 |
| | | 10000 | 10 | 0.36 | 0.16 | 0.20 | 0.14 | 0.15 | 0.10 | 0.08 | 0.07 | 0.07 | 0.04 |
| | | | 100 | 0.35 | 0.16 | 0.29 | 0.14 | 0.21 | 0.11 | 0.11 | 0.06 | 0.07 | 0.04 |
| | | 100000 | 100 | 0.75 | 0.13 | 0.71 | 0.14 | 0.64 | 0.17 | 0.54 | 0.17 | 0.31 | 0.19 |

Table S4: Performance in terms of AUROC to recover intra- (left) and intercellular (right) interactions from *in silico* tissues generated via mechanistic modeling for MISTy and SpaCeNet.

| AUROC (intracellular) | MISTy | SpaCeNet |
|---|---|---|
| Tissue 1 | 0.66 | 0.81 |
| Tissue 2 | 0.66 | 0.81 |
| Tissue 1+2 | 0.66 | 0.80 |

| AUROC (intercellular) | MISTy | SpaCeNet |
|---|---|---|
| Tissue 1 | 0.60 | 0.62 |
| Tissue 2 | 0.62 | 0.64 |
| Tissue 1+2 | 0.60 | 0.65 |

Table S5: Top 10 spatial interactions discovered by SpaCeNet on the StarMap data on natural scale (left), on log scale (middle) and for two copies of gene Mbp denoted as Mbp* on a natural scale (right).

| Recovered edges (natural scale) | | | Recovered edges (log scale) | | | Artificial Mbp2 colinear to Mbp (natural scale) | | |
|---|---|---|---|---|---|---|---|---|
| Gene 1 | Gene 2 | $\boldsymbol{\Delta\rho}$ | Gene 1 | Gene 2 | $\boldsymbol{\Delta\rho}$ | | | |
| Mbp | Flt1 | 0.327 | Sst | Pvalb | 0.178 | Mbp* | Mbp** | -0.173 |
| Ctgf | Gja1 | 0.102 | Mbp | Flt1 | 0.150 | Mbp* | Flt1 | 0.165 |
| Ctgf | Pcp4 | 0.083 | Ctgf | Pcp4 | 0.115 | Flt1 | Mbp** | 0.165 |
| Sst | Reln | 0.077 | Ctgf | Gja1 | 0.114 | Ctgf | Gja1 | 0.110 |
| Slc17a7 | Egr1 | 0.075 | Npy | Sst | 0.111 | Ctgf | Pcp4 | 0.083 |
| Cux2 | Gja1 | 0.060 | Sst | Vip | 0.107 | Sst | Reln | 0.078 |
| Cux2 | Egr1 | 0.055 | Cux2 | Pcp4 | -0.104 | Slc17a7 | Egr1 | 0.077 |
| Npy | Vip | 0.054 | Cux2 | Plcxd2 | 0.098 | Cux2 | Gja1 | 0.062 |
| Egr1 | Egr2 | 0.052 | Mgp | Cck | 0.098 | Cux2 | Egr1 | 0.057 |
| Cux2 | Pcp4 | -0.048 | Slc17a7 | Pcp4 | 0.095 | Npy | Vip | 0.055 |

Table S6: Top 10 spatial interactions discovered by SpaCeNet on the Drosophila data obtained on natural scale (left) and on log-scale (right)

| Natural scale | | | Log scale | | |
|---|---|---|---|---|---|
| Gene 1 | Gene 2 | $\boldsymbol{\Delta\rho}$ | Gene 1 | Gene 2 | $\boldsymbol{\Delta\rho}$ |
| sna | twi | -0.0360 | sna | twi | -0.0316 |
| ems | noc | 0.0301 | dan | danr | -0.0316 |
| Dfd | lok | 0.0210 | ems | noc | 0.0312 |
| Ance | CG10479 | 0.0205 | Ance | CG10479 | 0.0215 |
| dan | danr | -0.0193 | Dfd | lok | 0.0194 |
| cnc | erm | 0.0175 | apt | tll | -0.0159 |
| apt | tll | -0.0171 | cnc | erm | 0.0150 |
| Blimp-1 | brk | 0.0156 | Blimp-1 | brk | 0.0129 |
| CG14427 | kni | 0.0122 | kni | Nek2 | 0.0126 |
| cnc | kni | 0.0117 | cnc | kni | 0.0118 |

Table S7: Top 10 absolute highest values of $\Delta\rho$ for genes appearing in at least 30% of all cells (left) and genes appearing in 10% of cells (right).

| | 30% | | | 10% | |
|---|---|---|---|---|---|
| Gene 1 | Gene 2 | $\boldsymbol{\Delta\rho}$ | Gene 1 | Gene 2 | $\boldsymbol{\Delta\rho}$ |
| Mt2 | Mt1 | −0.363 | Mobp | Mbp | −0.448 |
| Mobp | Mbp | −0.355 | Sst | Npy | 0.374 |
| Ptgds | Apoe | 0.319 | Fth1 | Mbp | −0.356 |
| Il31ra | Camk1d | −0.287 | Mt2 | Mt1 | −0.347 |
| Bc1 | Hps5 | −0.287 | Il31ra | Camk1d | −0.331 |
| Gm42418 | Il31ra | 0.260 | Bc1 | Hps5 | −0.318 |
| Fth1 | Mbp | −0.218 | Mbp | Plekhb1 | −0.308 |
| Bc1 | Ppm1e | −0.208 | Nefl | Nefm | −0.296 |
| Olfm1 | Camk2n1 | −0.206 | Nrsn1 | Rgs4 | −0.257 |
| Plp1 | Cd81 | −0.200 | Bc1 | Ppm1e | −0.249 |

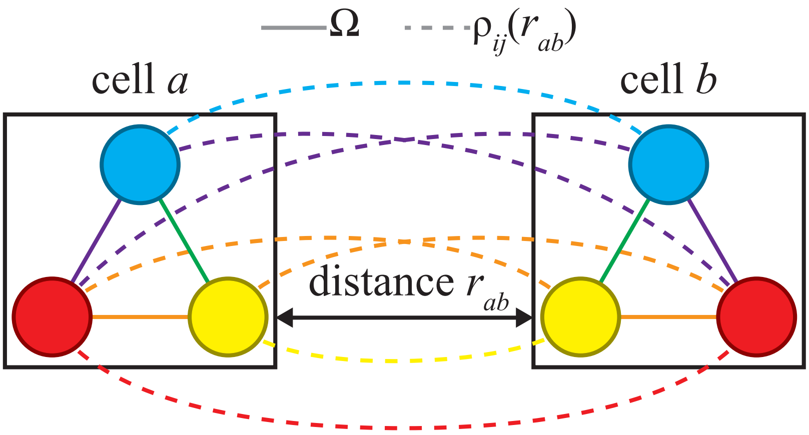


Figure S1: **Associations estimated by SpaCeNet:** SpaCeNet estimates intracellular and intercellular Spatial Conditional Independence (SCI) relationships reflected by network edges. Two cells $a$ (left) and $b$ (right) are shown. The circles of the same color represent the same molecular variable in each of the two cells. The solid lines represent edges of intracellular networks, indicating a direct association between molecular variables observed within the same cell. The dashed lines represent the intercellular network edges between cells $a$ and $b$, indicating a direct spatial association between respective molecular variables. The potentials $\rho_{ij}(r_{ab})$ parameterize the intercellular interaction strength between variables $i$ and $j$ between cell $a$ and cell $b$ at distance $r_{ab}$. For instance, the missing dashed green lines between the blue and yellow variable imply SCI between the two ($\rho_{ij}(r_{ab}) = 0$), while SCI does not apply between the yellow and red variable ($\rho_{ij}(r_{ab}) \neq 0$).

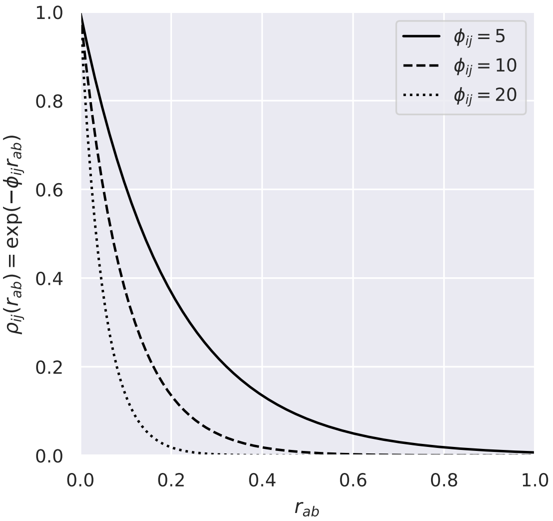


Figure S2: Illustration of different exponential potentials $\rho_{ij}(r_{ab}) = \exp(-\phi_{ij} r_{ab})$ used in the simulation studies, where $\phi_{ij} \sim \text{Unif}(5, 20)$ controls the interaction range. Small values correspond to long range associations ($\phi_{ij} = 5$, solid line), large values to short range associations ($\phi_{ij} = 20$, dotted line) and values in between to medium range associations ($\phi_{ij} = 10$, dashed line).

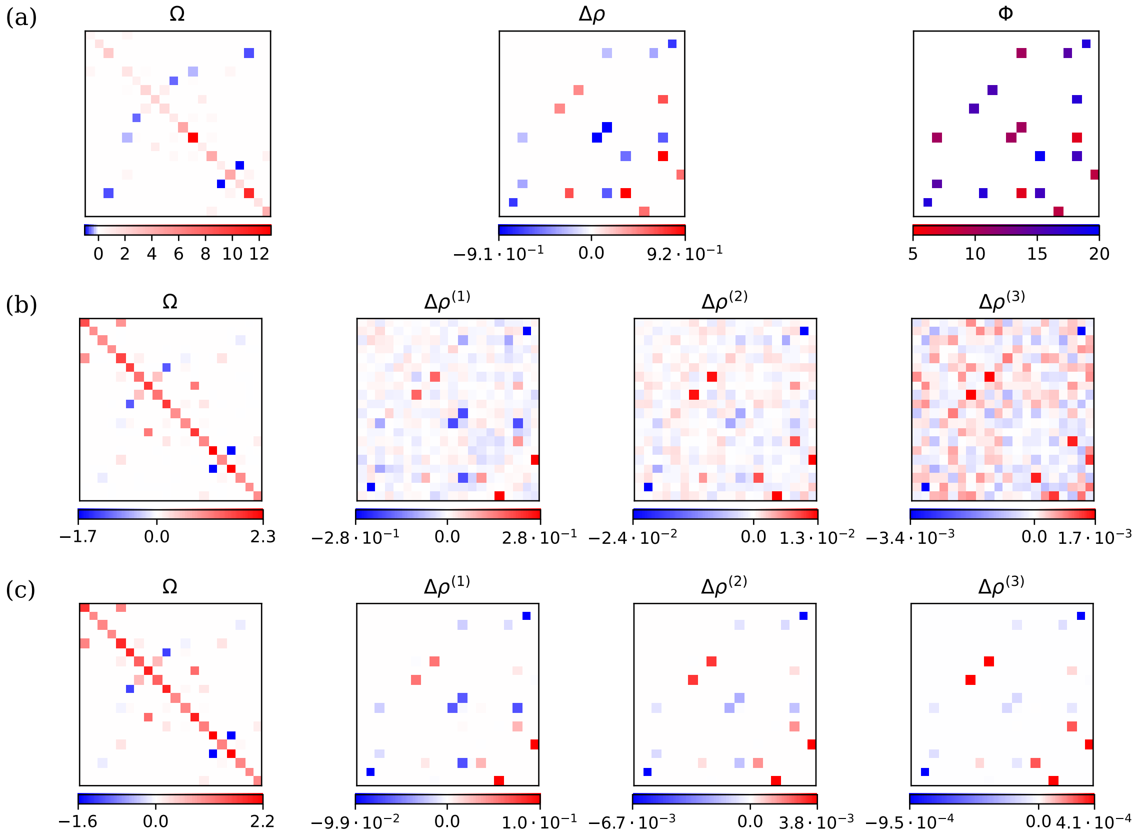


Figure S3: Parameters for a simulated data set with $n = 10^3,\ S = 100,\ \phi_{ij} \in [5, 20]$. (a) True parameters that have been used for sampling. (b) Estimated parameters, where the hyper-parameters were selected based on test-set loss, and (c) estimated parameters, where the hyper-parameters were manually chosen.

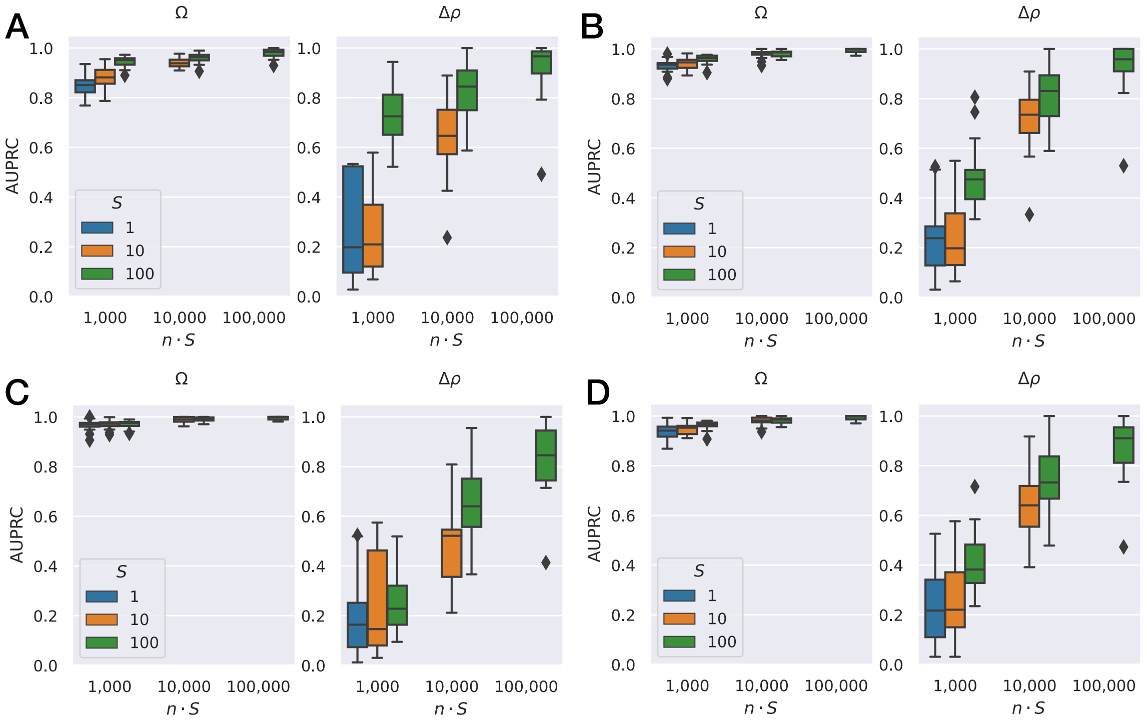


Figure S4: Edge recovery by SpaCeNet assessed in a simulation study using (A-C) fixed, radially decreasing potentials $\rho(r_{ab}) = \Delta\rho_{ij} \exp(-\phi_{ij} r_{ab})$ for all cell-cell interactions with constant range parameters $\phi_{ij} \in \{5, 10, 20\}$, respectively, and (D) flexible potentials $\rho_{ij}(r_{ab}) = \Delta\rho_{ij} \exp(-\phi_{ij} r_{ab})$ with $\phi_{ij} \sim \text{Unif}(5, 20)$ that mediate the interaction between molecular variables $i$ and $j$. The $y$-axes give the performance in terms of the area under the precision recall curve (AUPRC). Left figures correspond to the inner-cellular networks (the intracellular precision matrix $\boldsymbol{\Omega}$) and the right figures to the extracellular networks (the cell-cell interaction parameters $\boldsymbol{\Delta\rho}$). The $x$-axis stratifies the analysis with respect to total cell numbers $n \cdot S$, where $n$ is the number of cells within a measurement and $S$ the number of measurements. Here, $S = 1$ is shown in blue, $S = 10$ in orange, and $S = 100$ in green.

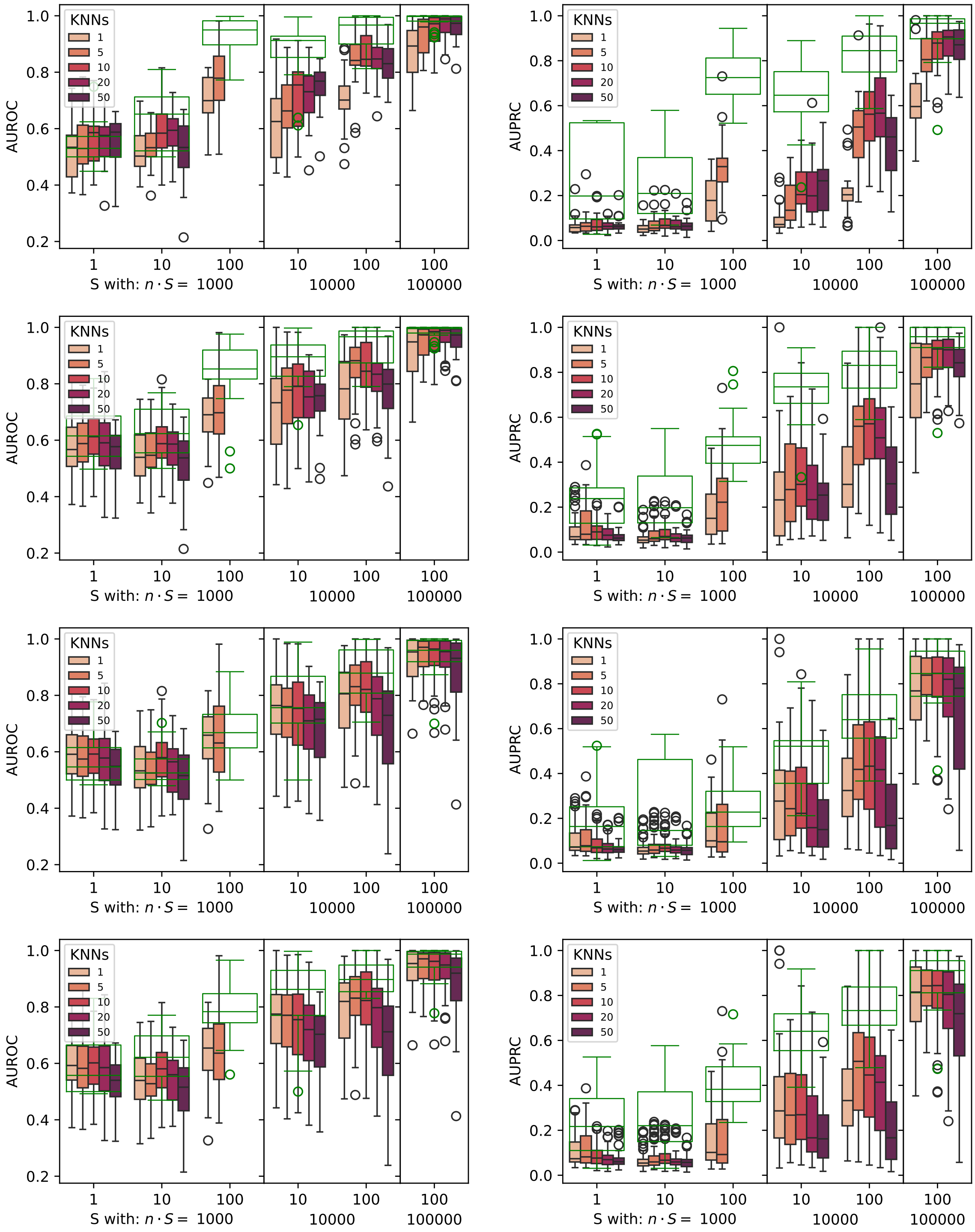


Figure S5: Edge recovery for intercellular associations $\Delta\rho$ of the correlation baseline considering a spatial environment given by the $K$ nearest neighbours. Row 1 to 4 correspond to simulation settings A-D, respectively (see Fig. S4). Results from SpaCeNet are shown in green.

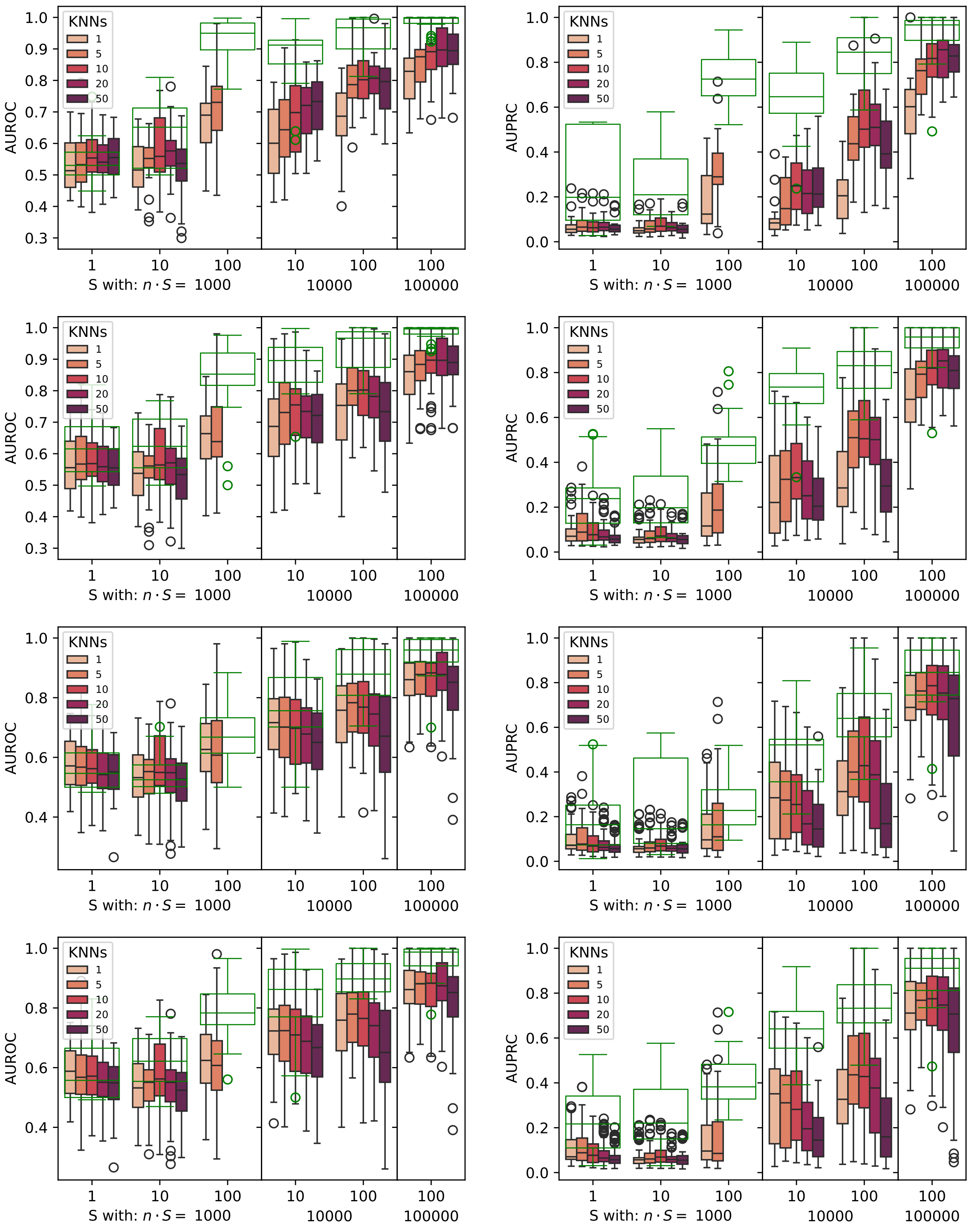


Figure S6: Edge recovery for intercellular associations $\Delta\rho$ of the partial correlation baseline considering a spatial environment given by the $K$ nearest neighbours. Row 1 to 4 correspond to Simulation settings A-D, respectively (see Fig. S4). Results from SpaCeNet are shown in green.

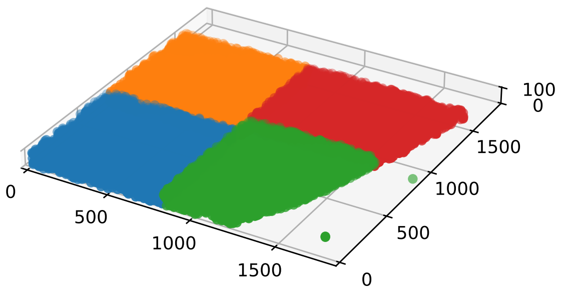


Figure S7: Cellular positions in the mouse visual cortex. Colors indicate the training (blue, orange, green) and validation (red) batches used for hyper-parameter screening and model development.

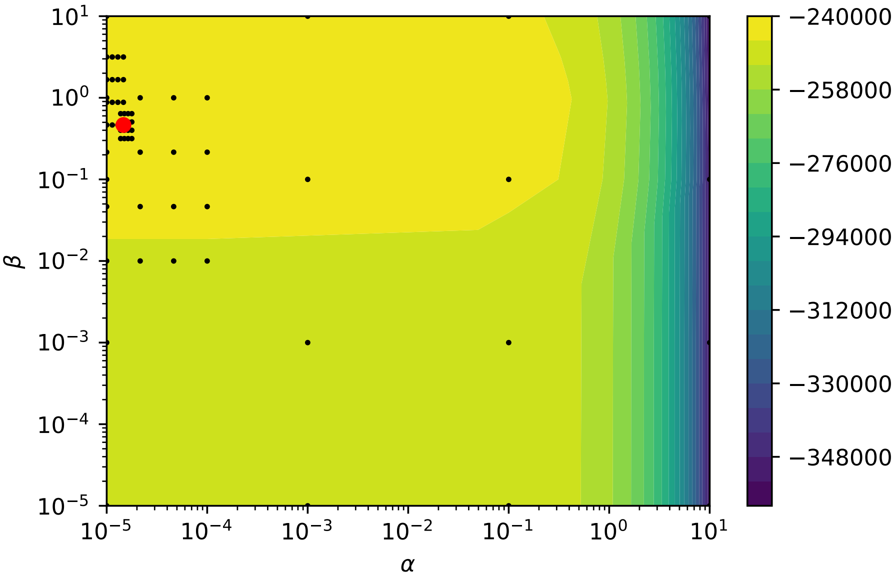


Figure S8: Evaluated hyper-parameter space and corresponding validation pseudo-log-likelihoods colored in blue (low values) to yellow (high values) based on the mouse visual cortex data provided by [Wang et al., 2018]. Black dots are tested hyper-parameters and the red dot corresponds to the optimal set of hyper-parameters in the grid search.

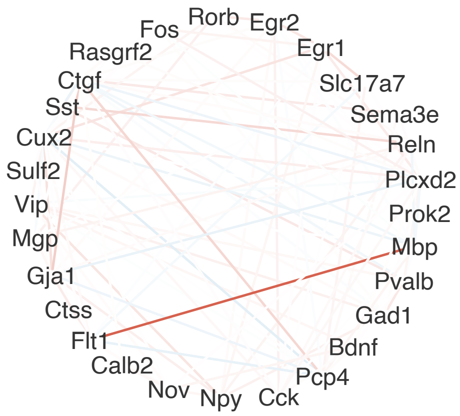


Figure S9: Network of spatial interactions between molecular variables estimated by SpaCeNet using data of the mouse visual cortex provided by [Wang et al., 2018]. Blue edges correspond to positive associations (negative entries of $\boldsymbol{\Delta\rho}^{(\cdot)}$) and red edges to negative associations (positive entries of $\boldsymbol{\Delta\rho}^{(\cdot)}$).

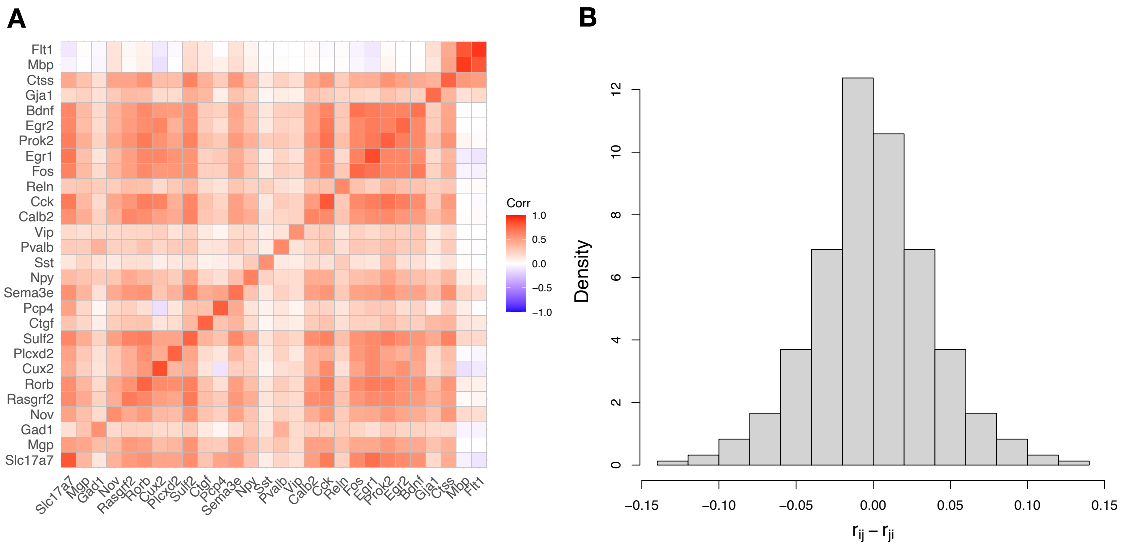


Figure S10: Figure A visualizes the Estimated pair-wise spatial correlations. The shown matrix is not symmetric by definition, although empirically approximately symmetric. This finding is supported by the histogram of differences $r_{ij} - r_{ji}$ (Figure B).

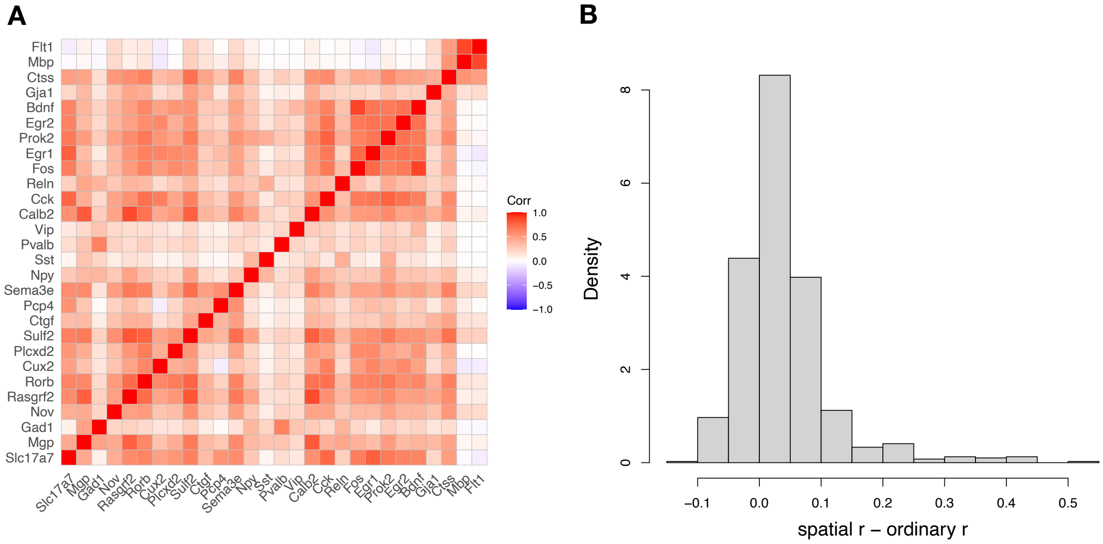


Figure S11: Figure A visualizes ordinary correlations calculated between gene expression levels based on the individual cells' molecular profiles. Figure B shows a histogram of the differences between the ordinary and spatial correlations.

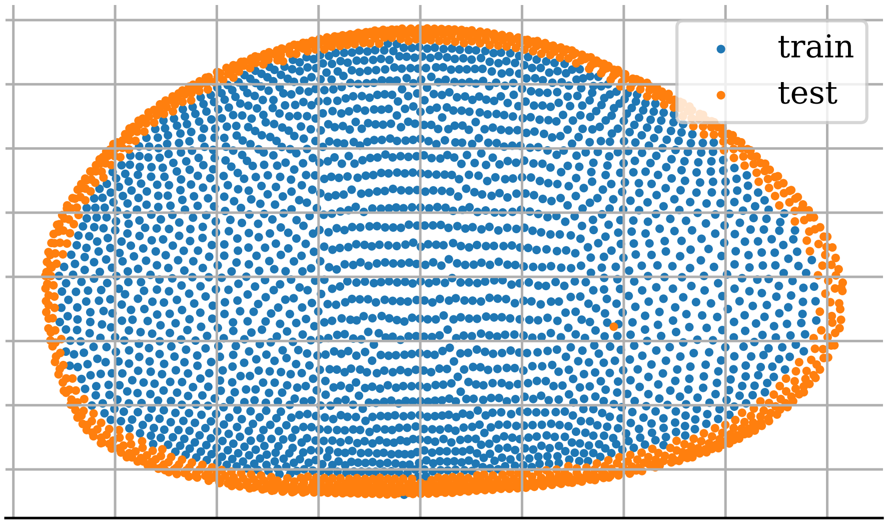


Figure S12: Spatial coordinates (mapped from 3D to 2D via a principal component analysis) of the virtual *Drosophila* embryo [Fowlkes et al., 2008] and the corresponding training/test splitting used for the SpaCeNet hyper-parameter screening shown in blue/orange, respectively.

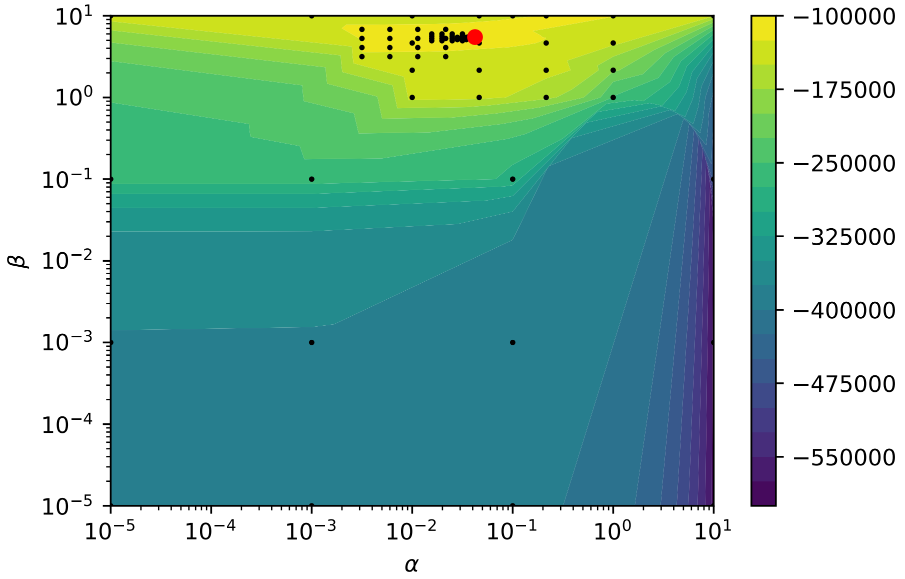


Figure S13: Evaluated hyper-parameter space and corresponding validation pseudo-log-likelihoods colored in blue (low values) to yellow (high values) based on the virtual *Drosophila* embryo data provided by [Fowlkes et al., 2008]. Black dots are tested hyper-parameters and the red dot corresponds to the optimal set of hyper-parameters in the grid search.

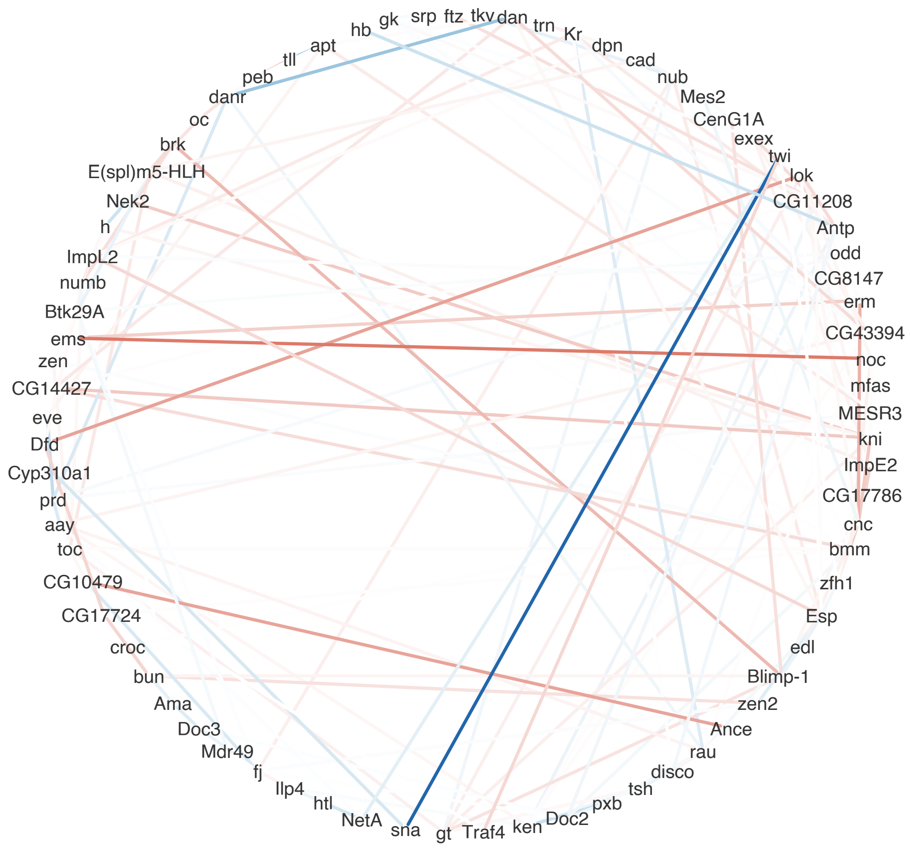


Figure S14: Network of spatial interactions between molecular variables estimated by SpaCeNet using data of the virtual *Drosophila* embryo provided by [Fowlkes et al., 2008]. Blue edges correspond to positive associations (negative entries of $\boldsymbol{\Delta\rho}^{(\cdot)}$) and red edges to negative associations (positive entries of $\boldsymbol{\Delta\rho}^{(\cdot)}$).

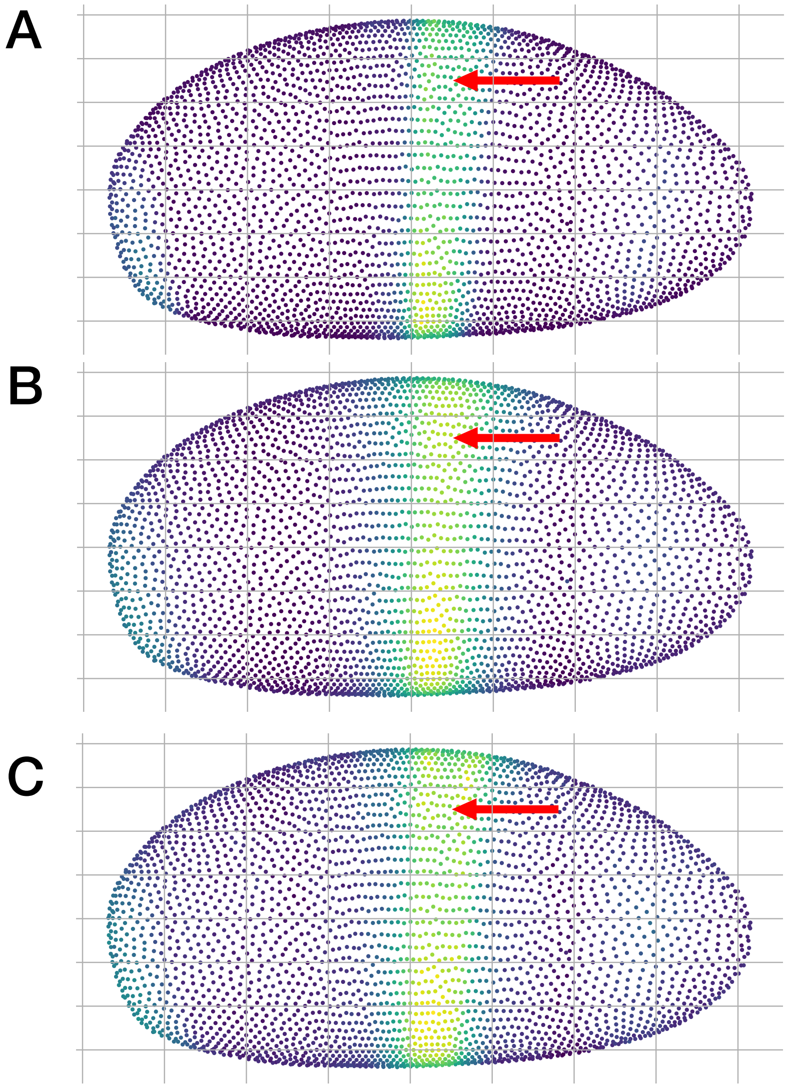


Figure S15: **SpaCeNet is an inferential tool which can predict gene expression from cellular context.** Figure A shows the expression of the Krüppel protein gene *Kr* based on the *Drosophila* blastoderm data of [Fowlkes et al., 2008]. Figure B shows the corresponding SpaCeNet prediction, where each cell's expression was predicted in a leave-one-cell-out approach (here, the density $f(\mathbf{x}^a|\mathbf{R}, \mathbf{X}^{\setminus a})$ was used). Figure C shows corresponding predictions based on cellular context and the expression of the remaining genes in the predicted cell (using density $f(x^a_j|\mathbf{R}, \mathbf{X}^{\setminus a}, \mathbf{x}^a_{\setminus j})$), meaning that the expression $\mathbf{x}^a_j$ is predicted using the expression levels of all other cells $\mathbf{x}^b$ with $b \neq a$ and the expression levels of cell $a$ except $j$. The red arrow highlights an area where the predictions of B and C differ most.

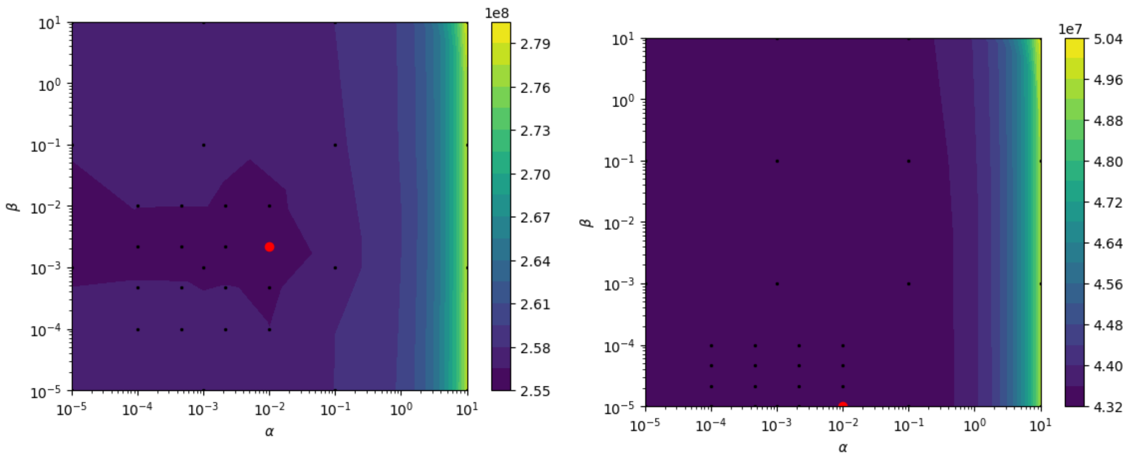


Figure S16: Evaluated hyper-parameter space and corresponding validation AIC colored in blue (low values) to yellow (high values) bassed on the MOSTA data, for genes appearing in at least 30% of all cells (left) and genes appearing in 10% of cells (right). Black dots are tested hyper-parameters and the red dot corresponds to the optimal set of hyper-parameters in the grid search.

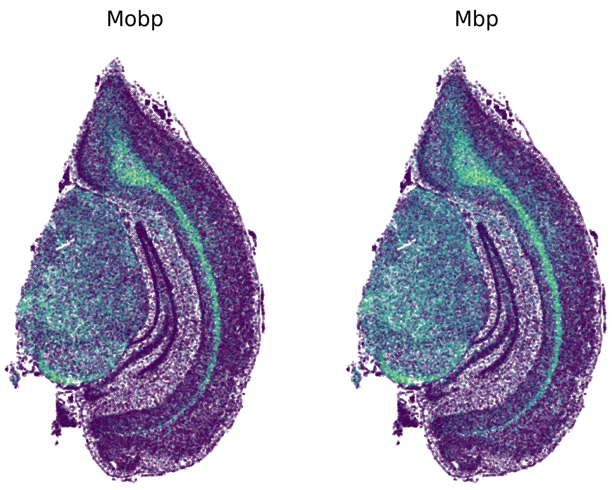


Figure S17: Scatter plot of the *Mobp* and *Mbp* genes in the MOSTA mouse adult brain colored in blue (low UMI count) to yellow (high UMI count).

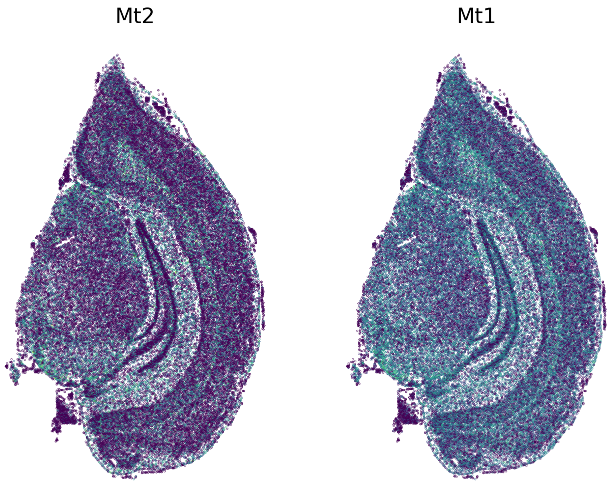


Figure S18: Scatter plot of the *Mt2* and *Mt1* genes in the MOSTA mouse adult brain colored in blue (low UMI count) to yellow (high UMI count).
